# The *glp-1* 3ʹ untranslated region regulates germline proliferation and promotes reproductive fecundity through multiple mechanisms

**DOI:** 10.64898/2026.03.23.713686

**Authors:** Peren Coskun, Sean P. Ryder

## Abstract

Germline development and successful embryogenesis depend upon the post-transcriptional regulation of maternal mRNAs. In *Caenorhabditis elegans*, the Notch-like receptor *glp-1* is necessary for germline progenitor cell proliferation in adults and anterior cell fate determination in embryos. The spatiotemporal patterning of GLP-1 protein has long served as a paradigm of maternal mRNA regulation in metazoans. The *glp-1* 3ʹUTR has been shown to be sufficient to pattern the expression of reporter genes, and multiple regulatory regions and RNA-binding protein interaction sites have been mapped. The RNA-binding proteins POS-1 and GLD-1 directly regulate *glp-1* mRNA via sequence specific interactions with motifs found in the *glp-1* 3ʹUTR. The impact of mutating the endogenous *glp-1* 3ʹUTR has not been studied, and the mechanism by which POS-1 and GLD-1 mediate repression is not understood. Here, we investigate the post-transcriptional mechanisms that govern *glp-1* expression, revealing that GLD-1 and POS-1 regulate this pattern through different pathways requiring different co-factors. Remarkably, mutations in the endogenous locus that disrupt either POS-1 or GLD-1 binding to the *glp-1* 3ʹUTR have minimal impact on reproductive fecundity. By contrast, a larger deletion that eliminates the binding of both has a strong effect on brood size, hatch rate, and displays an increase in the length of the germline mitotic region that corresponds with enhanced mitotic activity. Together, our results show that multiple post-transcriptional mechanisms work in concert to ensure robust GLP-1 patterning and thus maximize reproductive outcomes.

## INTRODUCTION

Female gametes are larger than their male counterparts and provide most of the cytoplasm to a newly fertilized zygote. The cytoplasm is loaded with mRNAs, proteins, and nutrients that are necessary for the zygote to progress through several rounds of cell division. In most species, early developmental decisions such as axis polarization, germline / soma segregation, and initial cell fate patterning decisions occur prior to zygotic gene activation (ZGA)—the point where zygotic transcription begins and the maternal mRNA supply is turned over [1]. As such, post-transcriptional regulatory mechanisms are thought to be crucial to regulating maternal mRNAs, ensuring that key cell fate specifying proteins are produced at the right place and time, thus enabling successful embryogenesis [2–6]. In the nematode model organism *Caenorhabditis elegans*, zygotic transcription begins at the four-cell stage in somatic lineages and after the >100 cell stage in the germline lineage [7–9]. Accordingly, several maternally supplied mRNAs have been shown to be important for early developmental decisions [10–14]. Mutations in these genes are often maternal effect lethal or maternal effect sterile [15].

In worms, the 3ʹUTR is the primary region that governs maternal RNA regulation in the germline [16]. Many *C. elegans* mRNAs go through a *trans* splicing reaction to add the 5ʹ cap structure, substituting the endogenous 5ʹ end of the transcript with one of two splice leader sequences [17]. As such, many transcripts are identical at the 5ʹ end. Reporter transgenes that fused the 3ʹUTRs of several germline transcripts to a fluorescence reporter gene and a generic germline promoter shows that the 3ʹUTR match the pattern of the endogenous gene in nearly all cases [16]. The sole exceptions were sperm-specific genes, whose patterning is governed by their promoters [16]. In a seminal discovery, the 3ʹUTR of *glp-1* mRNA was shown to be sufficient for the spatiotemporal patterning of a series of injected mRNA reporters, demonstrating patterning sufficiency in the absence of transcription, and revealing the post-transcriptional basis for GLP-1 regulation [4].

The *glp-1* gene encodes a Notch protein that receives cell signaling cues to activate promitotic transcriptional programs in the distal end of the germline [18]. In germline progenitor cells, GLP-1 receives a signal from the Delta-like ligand LAG-2, expressed on the somatic distal tip cell [19]. This interaction drives a conformational change in GLP-1 that promotes site-specific cleavage inside germline progenitor cells, releasing a C-terminal domain that enters the nucleus to promote transcription of mitotic genes, thus enabling germline progenitor cell renewal [20]. GLP-1 also promotes anterior cell fate specification in the embryo by transmitting signals between anterior and posterior blastomeres [18, 21]. GLP-1 is expressed in anterior lineage cells of early embryos, where it receives a signal from a second Delta-like ligand APX-1 to promote pharyngeal cell fate specification [5, 22, 23]. As such, the precise spatial patterning of GLP-1 is essential to both germline proliferation and anterior development in embryos. Accordingly, a loss of function *glp-1* mutant depletes progenitor cells rapidly and becomes sterile [18]. Embryos produced from *glp-1* mutant mothers fail to form the pharynx and arrest during morphogenesis [18, 22].

Transcripts encoding *glp-1* are observed throughout the germline and in all cells of the early embryo. By contrast, GLP-1 protein expression is restricted to the mitotic zone in the germline and the anterior cells of early embryos [4, 24]. The expression pattern is reported to be controlled by several RNA-binding proteins including MEX-3, OMA-1, PUF-3, PUF-5/6/7, GLD-1, POS-1 and SPN-4 (**Fig. 1A, B**) [25–30]. In the germline, GLD-1 represses GLP-1 expression in the meiotic zone [29], while OMA-1/2, PUF-3, MEX-3, and PUF-5/6/7 repress expression in oocytes [26, 27, 31]. In the embryos, GLD-1 and POS-1 repress GLP-1 expression in early posterior lineage cells [29]; and MEX-5/6 and SPN-4 promote GLP-1 expression in early anterior lineage cells [30].

**Fig. 1.**
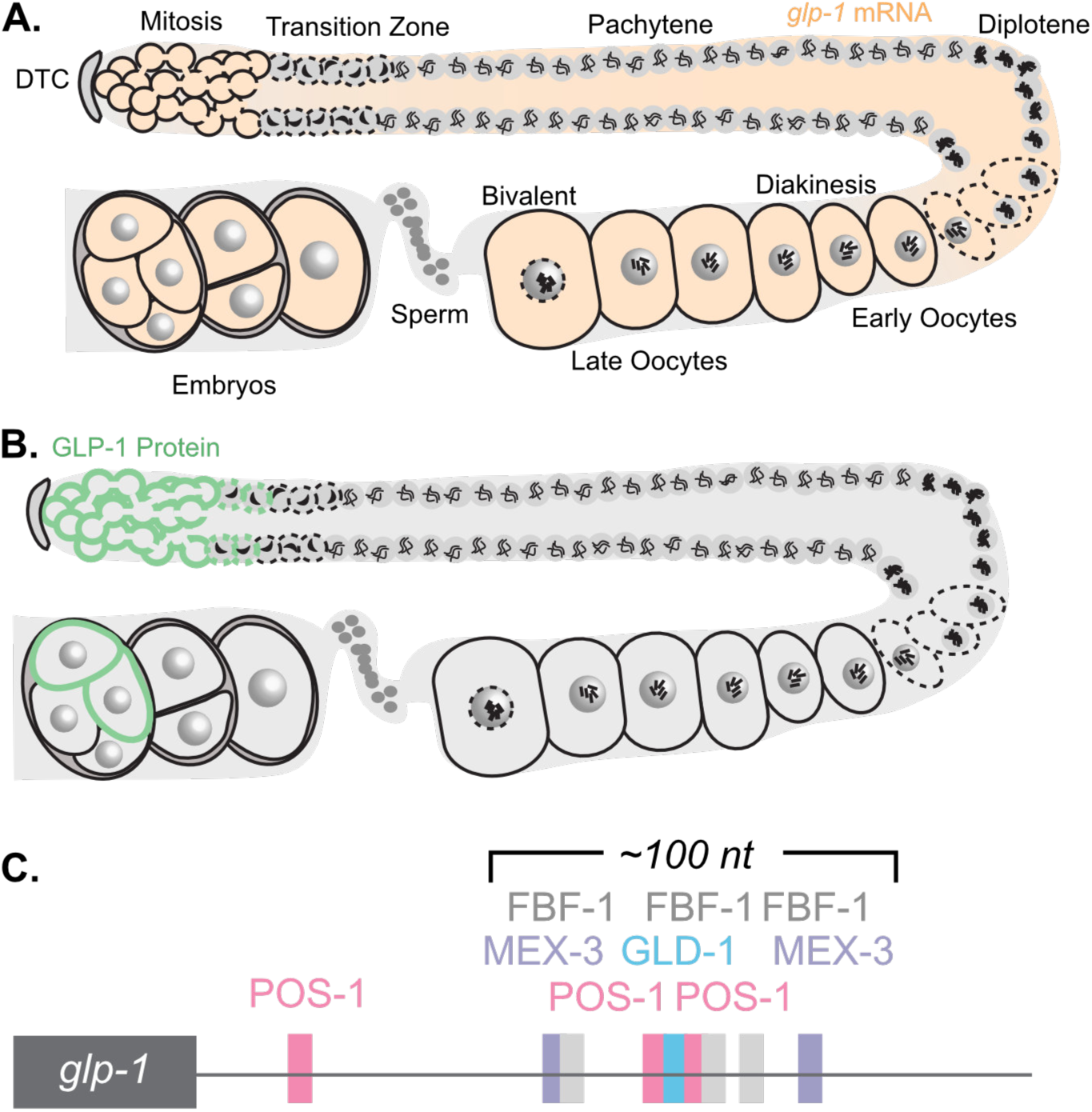
Expression pattern and 3ʹUTR diagram of *glp-1*. **A.** Anatomy of an adult hermaphrodite germline. The pattern of *glp-1* mRNA is shown in light orange. This image is modified from our previous figure published in [15] under the CC BY license. **B.** The pattern of GLP-1 protein is shown in the germline and early embryos in green. **C.** Diagram of the *glp-1* 3ʹUTR, with a map of both confirmed binding sites for POS-1 (pink) and GLD-1 (blue), as well as predicted sites matching motifs recognized by FBF (gray) or MEX-3 (purple).

Precise mutational studies of the *glp-1* 3ʹUTR in the context of injected mRNAs or transgenic reporters identified specific regions and precise binding motifs that are necessary for *glp-1* regulation (**Fig. 1C**) [4, 29, 32]. GLD-1 and POS-1, two key repressors of GLP-1 translation, recognize partially overlapping motifs within a region of the *glp-1* 3ʹUTR that governs its spatial control [29, 32]. GLD-1, a STAR/GSG/Maxi-KH domain RBP, binds to UACUC/AAY motifs (GLD-1 binding motif: GBM) [33–36], whereas POS-1—a CCCH type zinc finger protein—recognizes UAUUA/G(A/G/U)NNG motifs (POS-1 recognition element: PRE) [12, 37]. The motifs for both RBPs are present in a previously mapped *glp-1* repression element (GRE) [29]. A second POS-1 binding motif is positioned in the adjacent *glp-1* derepression element (GDE), a region previously shown to be necessary for reporter expression [29]. The physical interaction of each RBP to their respective element has confirmed in vitro. The GLD-1 binding site partially overlaps with both POS motifs, and *in vitro* binding studies showed the two RBPs compete for binding to this region of the *glp-1* 3ʹUTR [32].

Mutagenesis studies identified specific separation of function mutations that disrupt each binding site without perturbing binding to the other adjacent sites [32]. In transgenic reporter studies, the GLD-1 motif in the GRE and the POS-1 motif in the GDE were both required for repression in the posterior of embryos [32]. By contrast, the 5ʹ POS-1 motif appears to be non-functional in the reporter [32]. Mutation of the GLD-1 motif led to elevated expression in the meiotic zone of the adult germline, while mutation of the POS-1 motif led to reduced expression in the same region [32]. Although the *glp-1* 3ʹUTR has been studied for over three decades [4], there have been no mutational studies conducted at the endogenous locus, and the biological significance of the *glp-1* 3ʹUTR to the worm reproduction has yet to be characterized.

In this study, we combine genetic and molecular approaches to assess the mechanism of *glp-1* mRNA regulation and its biological impact on reproduction, focusing on the well-characterized GLD-1 and POS-1 binding elements. Our data reveal that GLD-1 and POS-1 repress *glp-1* through shortening of the polyA tail; however, our data also show that they appear to work via different pathways requiring different co-factors. Moreover, our work shows that translational control of *glp-1* by the cap-binding factor *ife-3* is important for repression in the germline, but neither the POS-1 nor GLD-1 binding sites contribute to this repression. Finally, our work shows that mutation of the POS-1 or GLD-1 binding sites in the endogenous *glp-1* 3ʹUTR has limited impact on reproductive fecundity in isolation, but deleting a region containing both sites has a strong reproductive phenotype—impacting fecundity, embryonic viability, and gene expression patterns.

## RESULTS

### The glp-1 3ʹUTR isoform distribution and polyA tail length are not impacted by loss of pos-1, gld-2, gld-3, mex-5, or mex-6 in embryos

The *glp-1* 3ʹUTR is necessary and sufficient to confer the GLP-1 expression pattern to a reporter transgene [4, 16, 32]. We set out to understand how this 3ʹUTR governs this pattern of expression. Transcripts can have different 3ʹUTRs as the consequence of alternative splicing (AS) or alternative polyadenylation (APA) [38]. Two separate studies mapped all *C. elegans* 3ʹUTRs using RNA collected from worms ranging in age from embryos to young adults [39, 40]. From these data, the *glp-1* 3ʹUTR was shown to have six possible isoforms that appear to derive from solely from APA. The UTR isoforms (A-F) range from 102 to 411 nucleotides in length (**Fig. 2A**). Isoforms A and B lack the previously mapped GRE and GDE elements, including the GLD-1 and POS-1 binding sites, but isoforms C to F do contain these motifs.

**Fig. 2.**
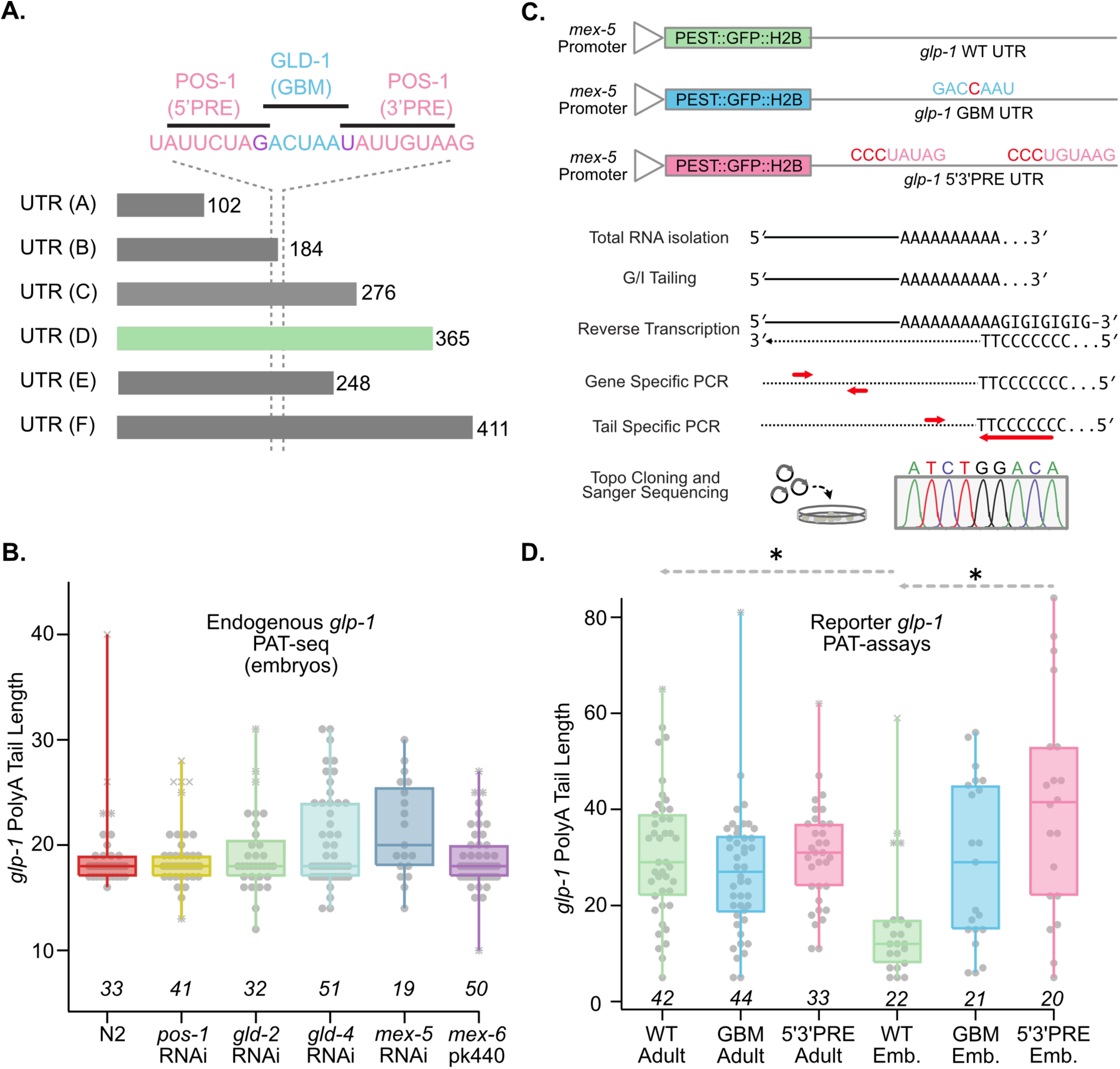
UTR isoform and polyA tail analysis of *glp-1* and *glp-1* 3ʹUTR reporter transcripts. **A.** The *glp-1* 3ʹUTR isoforms that can be formed by alternative polyadenylation are shown. We only observe isoform D (green) in our analysis of endogenous and reporter transcripts. The location of binding sites for POS-1 (pink) and GLD-1 (blue) is shown with dashed lines. **B.** Box plot of the polyA tail length distributions of endogenous *glp-1* polyA transcripts in embryos as a function of various RBP knock downs or mutants. The data are from genome wide PAT-test data published by Elewa et al. and available in Bioproject PRJNA248698 [41]. Each gray circle represents the polyA tail length from a *glp-1* read in the sequencing library (see methods). Asterisks represent near Tukey outliers, and “X” denotes far Tukey outliers (near outliers are 1.5X larger or smaller than the inner quartile range, far outliers are 3x). Whiskers represent the full range of the data. The number of reads analyzed is shown below. The horizontal line represents the median tail length. **C.** Diagram of the *glp-1* 3ʹUTR reporter transgenes used in this study (top) [32]. The mutations present in each strain are shown in red. The assay design for polyA tail length distribution measurements of the reporter transgenes is shown (bottom). Red arrows indicate gene and tail specific primer sets. The letter “I” indicates inosine. **D.** Box plot of the polyA tail distributions of the *glp-1* 3ʹUTR reporter transgenes from PAT assays outlined in (C). Each gray circle indicates the length of a polyA tail from an individual Sanger sequencing read. The number of colonies sequenced is shown below. The median, near and far outliers, and the full range of the data are shown as in (B). The color scheme for each reporter transgene variant follows the colors used in panel C. Dashed lines and black asterisks indicate statistically significant differences between the indicated distributions in a one-way ANOVA with Bonferroni multiple hypothesis correction (p_adj_ < 0.05).

To evaluate the distribution of *glp-1* 3ʹUTR isoforms in embryo samples, we analyzed reads corresponding to each isoform variant from published polyA-test sequencing (PAT-seq) data [41]. This data set contains samples obtained from N2 embryos, samples collected from embryos with *pos-1*, *gld-2*, *gld-3*, or *mex-5* RNAi treatment, as well as samples recovered from *mex-5(pk440)* mutant embryos. To recover reads mapping to the *glp-1* gene from each data set, raw sequencing data were downloaded from the NCBI sequence read archive corresponding to Bioproject PRJNA248658. The raw data were analyzed with custom scripts (see methods) containing a pattern matching algorithm to recover reads harboring each of the possible *glp-1* UTR isoforms, as well as the corresponding polyA tail length. The data show that only isoform D is present in embryo samples recovered from all genotypes and treatment conditions. The median polyA tail length is 18 adenosines for all samples except *mex-5* RNAi, where the median tail length is 20 adenosines (**Fig. 2B**). There is no statistically significant difference in the mean polyA tail length in all six samples (p_adj_ ∼ 1.0, **Supplemental Data Set 1**). The data do show that *glp-1* polyA tails in embryos are considerably shorter than the median polyA length in *C. elegans* (57 nucleotides) [42]. They are shorter than what is believed to be the minimal length necessary to support efficient translation (20 nucleotides) [42]. RNAi knockdown or mutation of five different RNA-binding proteins, including *pos-1*, does not appear to impact the median length or the isoform distribution in embryos.

### The glp-1 3ʹUTR polyA tail length is shorter in embryos than in young adults

To determine how polyA tail length is impacted by POS-1 and GLD-1 binding site mutations in the *glp-1* 3ʹUTR, we measured the polyA tail length distribution in our previously described *glp-1* 3ʹUTR reporter transgenes [37] (**Fig. 2C-D**). The advantage of measuring the distribution in the transgenic reporters is that RNAi of *pos-1*, *gld-2*, and *gld-3* is known to induce strong phenotypes in the germline or embryo [12, 43, 44]. These phenotypes could have indirect effects on the polyA tail length. The disadvantage is that endogenous *glp-1* transcripts are also present in these strains, which necessitates an assay that distinguishes reporter mRNAs from endogenous *glp-1* transcripts. Total RNA was recovered from adults and/or embryos from the WT *glp-1* 3ʹUTR reporter (WRM5: hereafter *glp-1* WT, **Table 1**), the GLD-1 binding motif mutant 3ʹUTR reporter (WRM6: hereafter *glp-1* GBM), or a reporter where both POS-1 binding sites are mutated (WRM9: hereafter *glp-1* 5ʹ3ʹPRE) [37]. Recovered transcripts were tailed with guanosine and inosine using established protocols [45] and then amplified using RT-PCR with a 5ʹ-polyC-TT-3ʹ tail primer and a forward primer that anneals to the GFP reporter, preventing endogenous transcript amplification (**Fig. 2C**). The amplicon was topo-cloned, transformed into *Escherichia coli*, and multiple individual colonies were sequenced for each sample using Sanger sequencing to estimate the isoform ratio and length of the polyA tails (**Fig. 2D**).

**TABLE 1:** Strains used in this study.

| Strain ID | Genotype | Approach | Reference |
| --- | --- | --- | --- |
| N2 | WT | N/A | [68] |
| WRM5 | *sprSi5[Pmex-5::MODC*<br>*PEST:GFP:H2B::glp-1 3'UTR cb-unc-*<br>*119(+)] II, unc-119(ed3) III* | mosSCI | [32] |
| WRM6 | *sprSi6[Pmex-5::MODC*<br>*PEST:GFP:H2B::glp-1 3'UTR(ΔGBM) cb-*<br>*unc-119(+)] II, unc-119(ed3) III* | mosSCI | [32] |
| WRM9 | *sprSi9[Pmex-5::MODC*<br>*PEST:GFP:H2B::glp-1 3'UTR(Δ5'3' PRE)*<br>*cb-unc-119(+)] II, unc-119(ed3) III* | mosSCI | [32] |
| JK5973 | *glp-1(q997[glp-1::2xOLLAS]) III* | CRISPR | [48] |
| WRM73 | *glp-1(spr16[q997\*(glp-1::2XOL-*<br>*LAS::Δ71_3'UTR]) III* | CRISPR | This paper |
| WRM83 | *glp-1(spr21[q997\*(glp-1::2XOL-*<br>*LAS::WT_3'UTR (HindIII site)]) III* | CRISPR | This paper |
| WRM84 | *glp-1(spr22[q997\*(glp-1::2XOLLAS::*<br>*5'3'PRE_3'UTR (HindIII site)]) III* | CRISPR | This paper |
| WRM109 | *glp-1(spr33[q997\*(glp-1::2XOL-*<br>*LAS::GBM_3'UTR (HindIII site)]) III* | CRISPR | This paper |
| EU552 | *glp-1(or178) III* | EMS | Un-<br>published |

Consistent with the genome-wide data, we only observe transcripts corresponding to isoform D. Our results show that the median polyA tail length is 29 in young adult *glp-1* WT animals, 27 in *glp-1* GBM, and 31 in *glp-1* 5ʹ3ʹPRE. The data demonstrates that mutation of the GLD-1 and POS-1 binding sites do not alter the polyA tail distribution in young adults (p_adj_ = 1.00, **Fig. 2D**). In embryo samples collected with the reporter transgenes, the polyA tail length is considerably shorter in *glp-1* WT embryos compared to young adults (*glp-1* WT_YA_ = 29, *glp-1* WT_emb_ = 12, p_adj_ = 3.8e-4, **Fig. 2D**), suggesting that polyA tail lengths are shortened between the adult germline and fertilized embryos. This observation is consistent with the genome wide study that showed short polyA tails on endogenous *glp-1* transcripts (**Fig 2B**). PolyA tail shortening is not observed in *glp-1* GBM or *glp-1* 5ʹ3ʹPRE reporters (*glp-1* GBM_YA_ =27, *glp-1* GBM_emb_ = 29, p_adj_ = 1; *glp-1* 5ʹ3ʹPRE _YA_ = 31, *glp-1* 5ʹ3ʹPRE _emb_ = 42, **Fig. 2D**). Together, the results suggest that derepression of the *glp-1* GBM and *glp-1* 5ʹ3ʹPRE reporter transgenes in embryos is caused in part by failure of a polyA tail shortening mechanism that acts on the wild-type 3ʹUTR.

### GLD-2 and GLD-4 cytoPAPs contribute to glp-1 polyA tail distribution in different ways

We were intrigued by the difference in embryonic polyA-tail distribution in the *glp-1* GBM and/or *glp-1* 5ʹ3ʹPRE reporter transgenes and wondered if cytoplasmic polyA polymerase activity contributes to derepression. GLD-2 and GLD-4 are both well-characterized cytoplasmic polyA polymerases found in early embryos [44, 46, 47]. We knocked *gld-2* or *gld-4* down using RNAi in all three reporter strains and quantified the GFP fluorescence intensity in each cell of dissected four-cell embryos, comparing the pattern and intensity of the fluorescence (**Fig. 3A**). In untreated *glp-1* WT embryos, reporter expression is biased towards anterior cells, as previously reported [32]. In untreated *glp-1* GBM and *glp-1* 5ʹ3ʹPRE mutant embryos, expression is increased and observed in all four cells (Fold Effect *glp-1* GBM / *glp-1* WT = 3.0; p_adj_ = <1e-12; *glp-1* 5ʹ3ʹPRE / *glp-1* WT = 3.4, p_adj_ = <1e-12, **Fig. 3A,B**) [32]. RNAi of *gld-2* or *gld-4* had no significant impact on the abundance or pattern of GFP fluorescence in *glp-1* WT UTR embryos (*glp-1* WT*_gld-2_* / *glp-1* WT_ctl_ = 1.0; p_adj_ = 1.0; *glp-1* WT*_gld-4_* / *glp-1* WT_ctl_ = 1.4, p_adj_ = 1.0; **Fig. 3C**). However, *gld-2* RNAi caused a decrease in the fluorescence intensity of both the *glp-1* GBM and *glp-1* 5ʹ3ʹPRE reporters (*glp-1* GBM*_gld-2_* / *glp-1* GBM_ctl_ = 0.73, p_adj_ = 0.00026; *glp-1* 5ʹ3ʹPRE*_gld-2_* / *glp-1* 5ʹ3ʹPRE_control_ = 0.72, p_adj_ = 6.47e-7; **Fig. 3D,E**). This suggests that GLD-2 contributes to the derepression that is observed when either the GLD-1 or POS-1 binding sites are mutated. Intriguingly, in *gld-4* RNAi animals, different results were observed in the GBM versus 5ʹ3ʹPRE reporters, with total fluorescence increasing in *glp-1* GBM but remaining unchanged in the 5ʹ3ʹPRE reporter (*glp-1* GBM*_gld-4_*/ *glp-1* GBM_ctl_ = 1.36, p_adj_ = 1.7e-7; *glp-1* 5ʹ3ʹPRE*_gld-4_* / *glp-1* 5ʹ3ʹPRE_ctl_ = 1.05, p_adj_ = 1.0; **Fig. 3D,E**). These results show that knock down of *gld-2* and *gld-4* have opposing effects on the *glp-1* GBM reporter, and that knock down of *gld-2*, but not *gld-4*, impacts the expression from the *glp-1* 5ʹ3ʹPRE reporter.

**Fig. 3.**
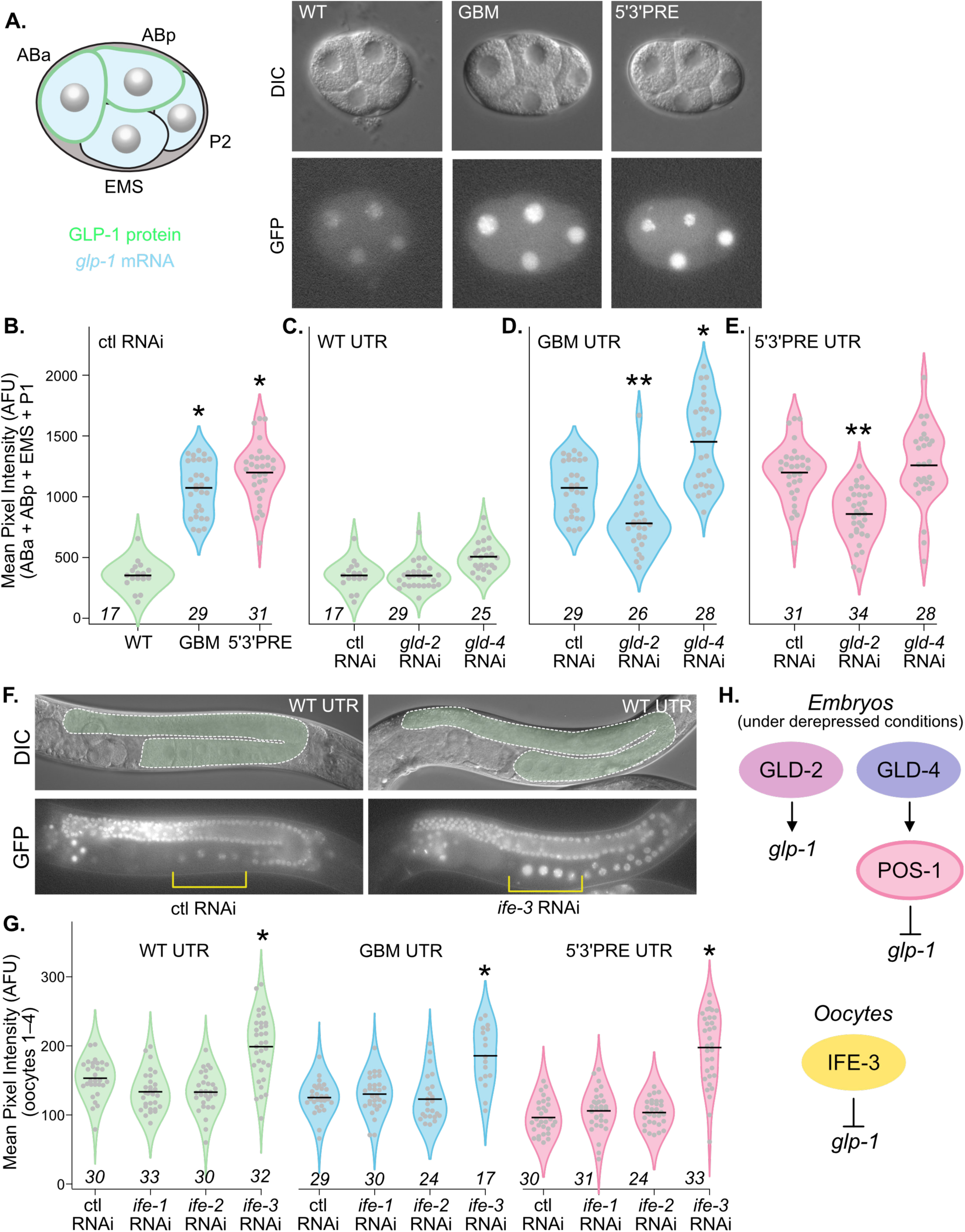
Impact of post-transcriptional regulatory mechanisms on *glp-1* 3ʹUTR reporter transgene expression. **A.** Diagram of a 4-cell embryo with the pattern of GLP-1 protein (green) and mRNA (light blue) shown (left). DIC (top) and GFP (bottom) images of 4-cell embryos recovered from each of the *glp-1* 3ʹUTR reporter transgenes [32]. **B.** Violin plots of the fluorescence intensity of each 4-cell embryo from each strain under control RNAi treatment. Each gray circle represents a unique embryo. The horizontal black line represents the mean. Asterisks indicate statistically significant increases in a one-way ANOVA with Bonferroni correction for multiple hypothesis testing (p_adj_ < 0.05). The color scheme for each variant follows figure 2C. The total number of 4-cell embryos imaged is shown below. **C.** Same as in B, except control (ctl), *gld-2*, and *gld-4* RNAi are used with the WT UTR reporter. **D.** Same as in C, except with the GBM mutant UTR reporter, and double asterisks indicate statistically significant decreases in a one-way ANOVA with multiple hypothesis correction. **E.** Same as in D except with the PRE mutant reporter. **F.** Images of young adult hermaphrodite germlines in the *glp-1* WT 3ʹUTR reporter transgenic line with both DIC optics (top) and GFP optics (bottom). In the DIC images, the germline is outlined in green. In the GFP images, the four most proximal oocytes are bracketed in yellow. Left is treated with control RNAi, right is treated with *ife-3* RNAi. **G.** Violin plot of the mean pixel intensity in the four most proximal oocytes in images of germlines from the WT UTR reporter strain (green), the GBM mutant UTR reporter strain (blue), and the PRE mutant UTR reporter strain (pink). The identity of the RNAi target for each experiment is shown. Each dot represents an individual germline. The black bar represents the median. The number of germlines imaged is shown below. The asterisks represent statical significance in a one-way ANOVA with Bonferroni correction. **H**. Model for *glp-1* activation by GLD-2, repression by GLD-4, and repression by IFE-3. Only repression by GLD-4 is strain-specific.

### Regulation of glp-1 3ʹUTR reporter transgenes by ife-3

We are also interested in understanding *glp-1* 3ʹUTR reporter transgene regulation in the germline. Several key regulatory factors, including translation initiation factors, cause sterility or embryonic lethality when mutated. As such, we chose to investigate their impacts on the *glp-1* 3ʹUTR reporter variants in oocytes. We chose three germline expressed eIF4E-binding proteins homologs (*ife-1*, *ife-2*, and *ife-3*) for RNAi knockdown. We measured the mean GFP intensity of the four most proximal oocytes in all three reporter strains treated with RNAi targeting each of these ten regulatory factors compared to controls. We observe no change in the mean fluorescence intensity of worms treated with *ife-1* or *ife-2* in any of the three strains (**Fig. 3F,G**, **Supplemental Data Set 1**). However, we observe a strong upregulation of all three reporters upon *ife-3* knockdown (Fold Effect *glp-1* WT*_ife-3_* / *glp-1* WT_ctl_ = 1.37, p_adj_ = 1.05e-9; Fold Effect *glp-1* GBM*_ife-3_* / *glp-1* GBM_ctl_ = 1.48, p_adj_ = 2.89e-8; Fold Effect *glp-1* 5ʹ3ʹPRE*_ife-3_* / *glp-1* 5ʹ3ʹPRE_ctl_ = 2.05, p_adj_ = <1e-7). Taken together, the data suggest that GLD-2 activates transgene expression in embryos, but only under derepressed conditions (**Fig. 3H**). In contrast, GLD-4 represses transgene expression in embryos, potentially via the POS-1 binding site. IFE-3 represses transgene expression in oocytes, but this mechanism seems to be independent of the POS-1 or GLD-1 binding sites.

### Mutation of the GLD-1 and POS-1 binding sites in the endogenous glp-1 gene

To understand the physiological impacts of mutating the GLD-1 and POS-1 binding sites in the endogenous *glp-1* locus, we used CRISPR-Cas9 mutagenesis to make precise mutations in an OLLAS-tagged *glp-1* strain corresponding to the variants we previously made in our reporter strains (**Fig. 4A**) [32, 48]. Our design also includes removal of a protospacer adjacent motif (PAM) to prevent re-cleavage of the knock in mutation, and mutation of a HindIII restriction site to facilitate screening. In total, we produced three strains: *glp-1(spr21[q977*])*, a control strain with the PAM site deletion and the HindIII site insertion that is otherwise wild-type (WRM83: hereafter *glp-1(WT*)*); *glp-1(spr33[q977*])* which is identical to *glp-1(WT*)* except it also harbors the GBM mutation (WRM109: hereafter *glp-1(GBM)*); and *(glp-1(spr22[q977*])* which harbors the 5ʹ3ʹPRE mutation (WRM84: hereafter *glp-1(5ʹ3ʹPRE)*, **Table 1**). All three strains can be easily propagated as homozygotes.

**Fig. 4.**
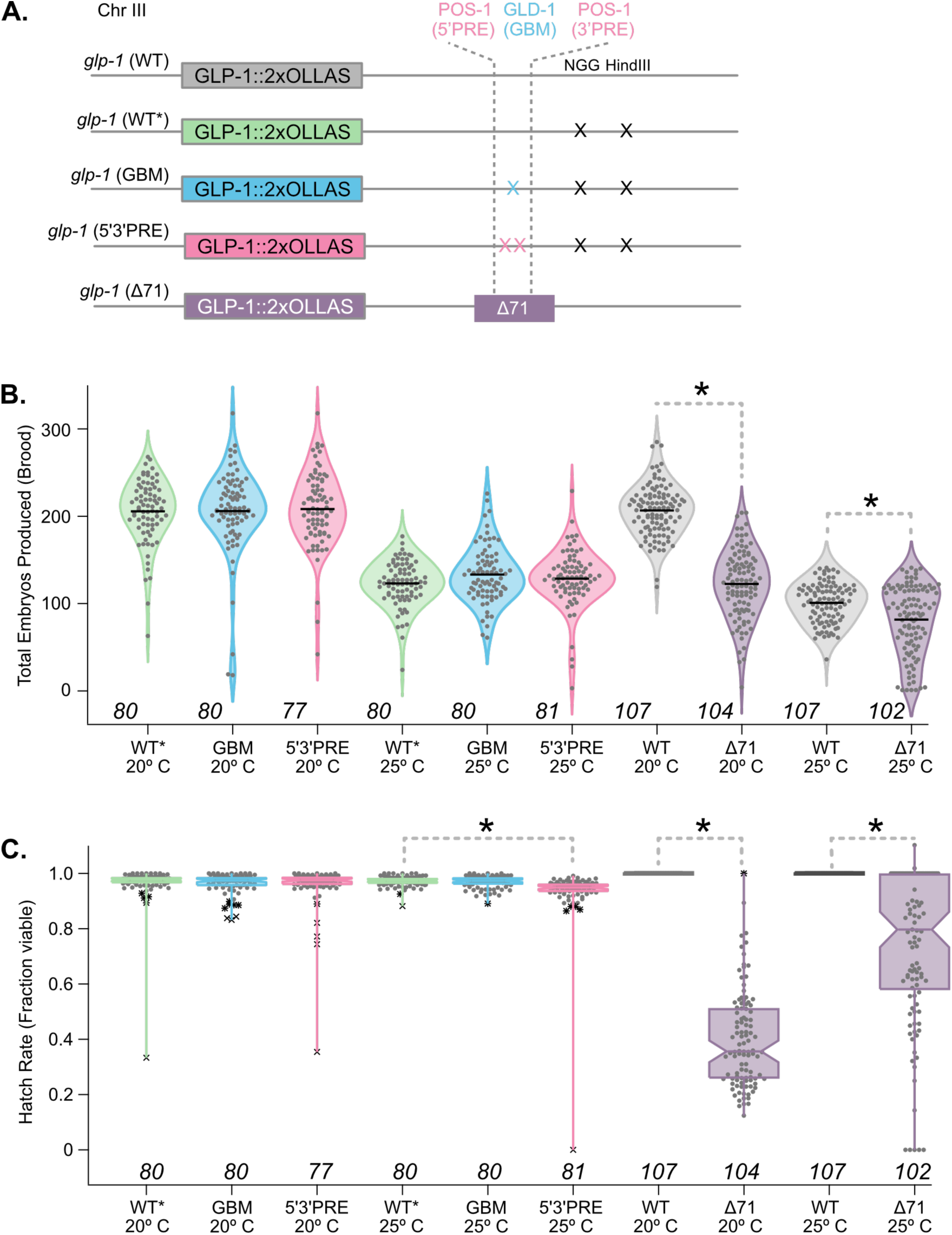
CRISPR *glp-1* 3ʹUTR alleles made in the endogenous *glp-1(q977[glp-1::2X OLLAS*]) genetic background [48]. The position of the GLD-1 (blue) and POS-1 (pink) binding sites are shown. The blue and pink “X” marks represent the same mutations as diagrammed in figure 2C in the context of the endogenous locus. The black “X” marks indicate deletion of the PAM and mutation of a HindIII site to simplify screening (aagctt–aagcGt). The purple box denoted Δ71 indicates the position of a 71 base pair deletion. The color of the GLP-1::2xOLLAS boxes corresponds to the quantitation shown in part B and C. **B.** Violin plots representing the distribution of the total number of embryos produced by a single hermaphrodite (each gray circle represents the total brood from one animal). The genotype, growth temperature (20°C and 25°C), and the number of animals assessed is shown per violin. Black bars represent the mean. The colors correspond to genotype as per (A). Asterisks indicate statistical significance in a one-way ANOVA with Bonferroni correction for multiple hypothesis testing. **C.** Box plots of the hatch rate of the embryos produced from each animal in panel (B). The individual gray circles are the hatch rate per animal, the asterisks and “X” marks represent near and far Tukey outliers as in figure 2B. The total number of animals assessed is listed below. The bold asterisks and gray dashed lines represent statistical significance in a one-way ANOVA with Bonferroni correction. The notches and horizontal lines indicate the mean.

To determine whether the GBM and 5ʹ3ʹPRE mutations affect reproductive fecundity, we measured the total brood size and hatch rate for all three strains. Under standard growth conditions at 20°C, the brood size and hatch rate are nearly identical for *glp-1 (WT*)*, *glp-1(GBM)*, and *glp-1 (5ʹ3ʹPRE)* alleles (**Fig. 4B,C**, **Supplementary Data Set 1**). Because 3ʹUTR mutations in other maternal transcripts have shown enhanced phenotypes at elevated temperatures [49, 50], we repeated the measurements with worms grown at 25°C. Under these conditions, the total brood size is similar in all three strains, but the hatch rate was mildly reduced in *glp-1(5ʹ3ʹPRE)* compared to the *glp-1(WT*)* allele (Fold effect: *glp-1(5ʹ3ʹPRE)* / *glp-1(WT*)* = 0.96, padj = 0.001, **Fig. 4B,C**). The hatch rate in *glp-1(GBM)* is unaffected. The results show that mutations that cause strong derepression of a *glp-1* reporter in embryos do not cause strong reproductive phenotypes when engineered into the endogenous locus, suggesting that reproduction is robust to dysregulation mediated by loss of either POS-1 or GLD-1 binding sites.

### Mutation of a region including both GLD-1 and POS-1 sites reduces fertility

Next, we used CRISPR to make a deletion of the *glp-1* 3ʹUTR (*glp-1(spr16[*q977])*) that removes both POS-1 binding sites, the GLD-1 binding site, and adjacent sites predicted to be recognized by FBF-1/2 and/or MEX-3. In total, this deletion precisely removes 71 base pairs from the 3ʹUTR (hereafter *glp-1(Δ71)*, **Fig. 4A**). In designing this deletion, we were able to identify a guide RNA that recognizes the region to be deleted in our intended mutation, obviating the need for the PAM deletion and HindIII site insertion used in making the precise binding site alleles described above. Despite deleting a region of the 3ʹUTR that has been shown to be sufficient for spatial patterning of a reporter [4], *glp-1(Δ71)* is viable and easily propagated as a homozygote. To assess the impact of this mutant on worm reproduction, we conducted brood size assays at both 20°C and 25°C. At 20°C, we observed a 1.7-fold reduction in the mean brood size of *glp-1(Δ71)* compared to the parent strain (*glp-1(q977)*), hereafter *glp-1(WT)*, p_adj_ = <1e-10, **Fig. 4B**). We also observed a 2.5-fold decrease in the hatch rate of *glp-1(Δ71)* embryos compared to the background strain (p_adj_ = <1e-10, **Fig. 4C**). In parallel to our findings at 20°C, we observed reduced brood size and hatch rate at 25°C for *glp-1(Δ71)* compared to the control (Fold effect reduction in total brood = 1.2, p_adj_ = 0.00003; Fold effect reduction in hatch rate = 1.35, p_adj_ = <1e-10, **Fig. 4B,C**). The difference in magnitude could be caused by the reduction of the brood size in the control *glp-1(WT)* strain at elevated temperature as opposed to temperature-dependent rescue in the mutant. The results show that the *glp-1* 3ʹUTR region containing the GLD-1, POS-1, and adjacent RBP-binding sites contributes to the total fecundity and successful embryogenesis, both under standard growth conditions and at elevated temperature.

### The Δ71 glp-1 3ʹUTR mutation produces terminal embryos that are distinct from a loss-of-function allele

On average, when incubated at 20°C, more than 60% of *glp-1(Δ71)* embryos fail to hatch. Loss-of-function *glp-1* mutants also produce embryos that fail to hatch, but with complete penetrance. We sought to compare the *glp-1(Δ71)* phenotype to the loss-of-function phenotype. We used DIC microscopy to image embryos harvested from *glp-1(WT)*, *glp-1(Δ71)*, and *glp-1(or178)*, which encodes a strong temperature sensitive loss-of-function allele (hereafter *glp-1(ts)*). Embryo images were binned by developmental stage and annotated based upon their appearance as normal or abnormal (**Fig. 5**). All early-stage embryos (one cell to 64-cell) produced by *glp-1(WT)* and *glp-1(ts)* looked normal in appearance (n=41, n=27, **Fig. 5A-C**). By contrast, 33% of *glp-1(Δ71)* early embryos appeared defective (n=9/27, p_adj_ = 1.4e-4, **Fig. 5B**). In later stage embryos (65-cell to pretzel stage), 90% of embryos were normal in *glp-1(ollas)* (n=26/29), while just 61% of *glp-1 (Δ71)* (n=64/105, p_adj_ = 0.007) and 52% of *glp-1(ts)* embryos (11/21, p_adj_ = 0.006) appeared normal (**Fig. 5D-F**). Most of the terminal *glp-1(ts)* embryos resembled the published *glp-1* loss-of-function phenotype with arrest after morphogenesis begins (**Fig. 5F**) [18]. In contrast, the *glp-1(Δ71)* embryos appeared to show a more variable phenotype, including regions of bundled cells and zones of clearing presumably due to apoptosis (**Fig. 5B,E**). These defects appear to manifest after the 16-cell stage but before morphogenesis. A quantitation of these results is shown in **figure 5G**. Together, our results suggest that the cause for embryonic lethality in *glp-1(Δ71)* embryos is distinct from loss of function *glp-1* alleles.

**Fig. 5.**
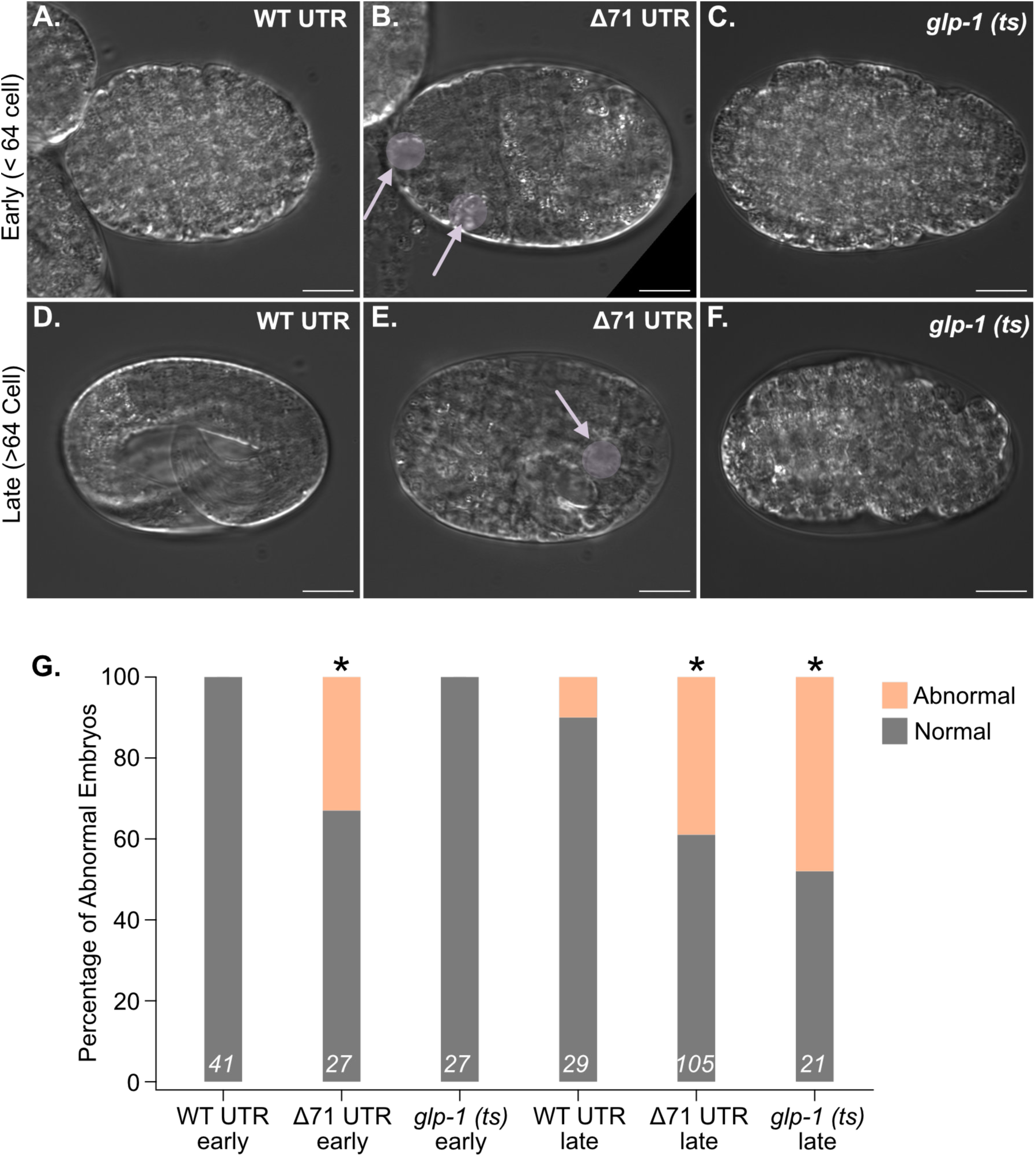
Terminal phenotype of *glp-1(Δ71) embryos*. **A.** DIC images of early and late embryos from the *glp-1(WT)*, *glp-1(Δ71)*, and *glp-1(ts)* strains. The arrows and circles identify zones of clearing presumably due to apoptosis that appear to be unique to the *glp-1(Δ71)* strain. Embryonic defects appear to manifest in younger embryos than in *glp-1(ts)* embryos. **B.** Stacked bar graph representing the percentage of normal vs. abnormal embryos observed in all three strains. The orange color reflects the fraction abnormal. Asterisks indicate statistical significance in a Chi Square test comparing each mutant to WT, with Bonferroni correction for multiple hypothesis testing.

### Increased germline mitosis in glp-1(Δ71) mutants

In the germline, GLP-1 protein expression is restricted to the distal end [24], and the repression of *glp-1* translation by the RNA-binding protein GLD-1 is thought to promote meiosis [29]. Consistent with this model, a gain-of-function *glp-1* allele forms a germline progenitor cell tumor [51], phenocopying strong loss-of-function *gld-1* alleles [10]. To evaluate the impact of the 3ʹUTR deletion on GLP-1 expression pattern in the germline, we dissected gonads from *glp-1(WT)* and *glp-1(Δ71)* young adult hermaphrodites, fixed them, and stained for the OLLAS tag on *glp-1*. We also imaged with DAPI to assess the morphology of the nuclei and determine the length of the mitotic zone (**Fig. 6A-F**). OLLAS immunofluorescence reveals that GLP-1 protein is expressed in both wildtype and Δ71 mutants in the distal arm of the germline to approximately the same level (**Fig. 6A**, **D**). To quantify the pattern of GLP-1 expression, we measured the distance from the distal tip of the gonad to the end of the OLLAS positive region in both *glp-1(WT)* and *glp-1(Δ71)* embryos. In gonads dissected from control animals, the average length of the OLLAS positive region is 113.2 ± 25 microns. In mutants, the length increases to 117.9 ± 20.7 microns, suggesting that GLP-1 expression is not expanded proximally in the Δ71 3ʹUTR deletion (p = 0.47, **Fig. 6G**).

**Fig. 6.**
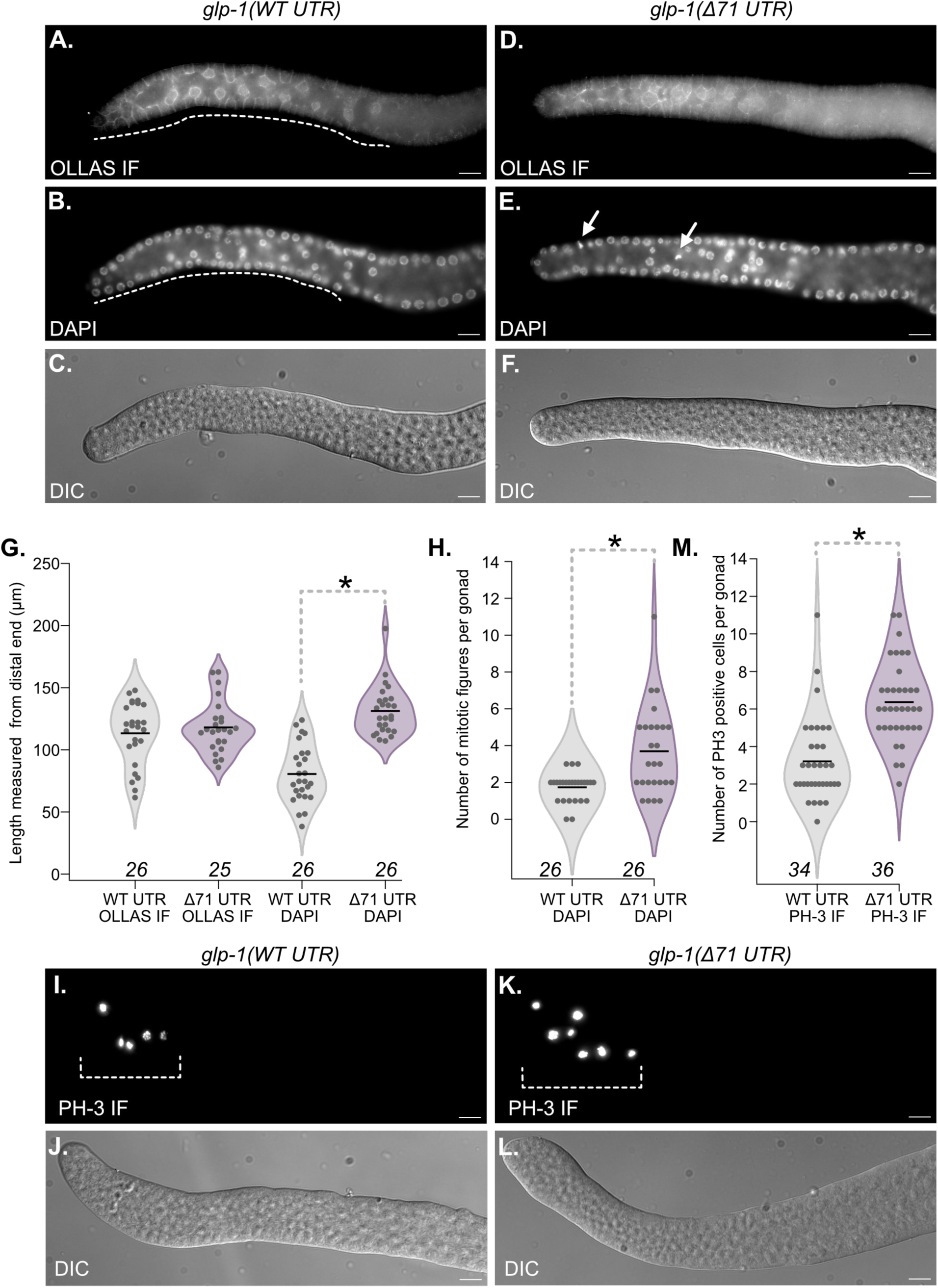
Increased mitosis in the distal germline of *glp-1(Δ71)* embryos. **A.** Anti-OLLAS immunofluorescence, **B.** DAPI staining, and **D.** DIC images of fixed gonads from *glp-1(WT)* worms. The white dashed line in (A) indicates an example of the distance of OLLAS staining for this gonad. The white dashed line in (B) indicates the distance from the distal end to the first meiotic nucleus. **D-F.** are images collected as in (A-C) except they are of fixed gonads from *glp-1(Δ71)* worms. The white arrows in panel E indicate mitotic figures. **G.** Violin plots representing the distribution of lengths of OLLAS staining (left) or the distribution of distance from the distal tip to the first meiotic nucleus (right) for both strains. The violin plots are colored to match the strains shown in figure 4A. Asterisks indicate statistical significance in a two-tailed t-test. The number of gonads analyzed is given below. **H.** Violin plots of the distribution of mitotic figures in the DAPI images in both strains. Coloring and statistical analysis are as in (G). **I.** Anti-PH-3 immunofluorescence and **J.** DIC images of fixed gonads dissected from *glp-1(WT)* animals. The white dashed bracket indicates PH-3 positive cells in the distal end of the germline. **K-L.** Anti-PH-3 and DIC images as in (I) and (J) except gonads were dissected from *glp-1(Δ71)* animals. **M.** Violin plot of the distribution of the number of PH-3 positive cells in both strains. Coloring and statistics are as in panel (G).

GLP-1 promotes mitotic gene expression in the distal end of the germline [18, 24]. To assess whether mitosis is increased in the 3ʹUTR mutant, we counted the number of distal mitotic figures present in the DAPI channel of the OLLAS immunofluorescence images described above. We also measured the overall length of the mitotic zone, defined as the distance from the distal end of the germline to the first crescent-shaped meiotic nucleus [52]. We observe an increase in the average number of mitotic figures in Δ71 mutants compared to controls (Δ71 = 3.69 ± 2.37, n = 26; control = 1.73 ± 0.77, n = 26; p = 0.00021, **Fig. 6H**). This corresponds to an increase in the average length of the mitotic zone in mutants (Δ71 = 131.2 ± 20, n = 26; control = 80.5 ± 23.2, n = 26; p = 3.7e-11, **Fig. 6G**). As an independent measure of increased mitosis, we stained for PH-3 positive nuclei in in the distal end (**Fig. 6I-L**). We observe twice as many PH-3 positive nuclei in the *glp-1(Δ71)* mutant compared to control (Mean PH-3+ (Δ71) = 6.4 ± 2, n = 34; PH-3+ (control) = 3.2 ± 2, n = 36, p = 6.6e-8, **Fig. 6M**). Together, the data show that the pattern of GLP-1 expression is enlarged in the Δ71 mutant, and this increase correlates to an increase in the number of germline progenitor cells undergoing mitosis.

### Mutation of the glp-1 3ʹUTR leads to changes in stress response and metabolism gene expression

To understand how dysregulation of *glp-1* expression in *glp-1(Δ71)* mutants impacts gene expression, we performed RNA-seq on age matched young adult hermaphrodites from synchronized populations of *glp-1(ollas)* and *glp-1(Δ71)* mutants. We also collected samples from synchronized young adult *glp-1(WT)*, *glp-1(GBM)*, and *glp-1(5ʹ3ʹPRE)* animals. Data were analyzed with the OneStopRNASeq pipeline [53], and gene expression differences between genotypes were assessed using DESeq2 with a log_2_ fold change cutoff of ≤ ±0.585 and a p_adj_ cutoff of ≤ 0.05 (**Fig. 7A-C**) [54]. The full analyses are available in **Supplemental Data Set 2**. Comparing *glp-1(ollas)* to *glp-1(Δ71)*, we observe 81 upregulated and 405 downregulated genes. Comparing *glp-1(WT)* to *glp-1(GBM)* and *glp-1(5ʹ3ʹPRE)* mutants, we see far fewer gene expression changes. We observe seven upregulated and one downregulated in *glp-1(GBM)*, and nine upregulated with six downregulated in *glp-1(5ʹ3ʹPRE).* Five of the 81 upregulated genes in *glp-1(Δ71)* are also upregulated in the *glp-1(GBM)* or *glp-1(5ʹ3ʹPRE)* mutants (**Fig. 7D**). Just one of the 405 downregulated genes in *glp-1(Δ71)* is shared with another mutant of the *glp-1* 3ʹUTR (5ʹ3ʹPRE, **Fig. 7E**).

**Fig. 7.**
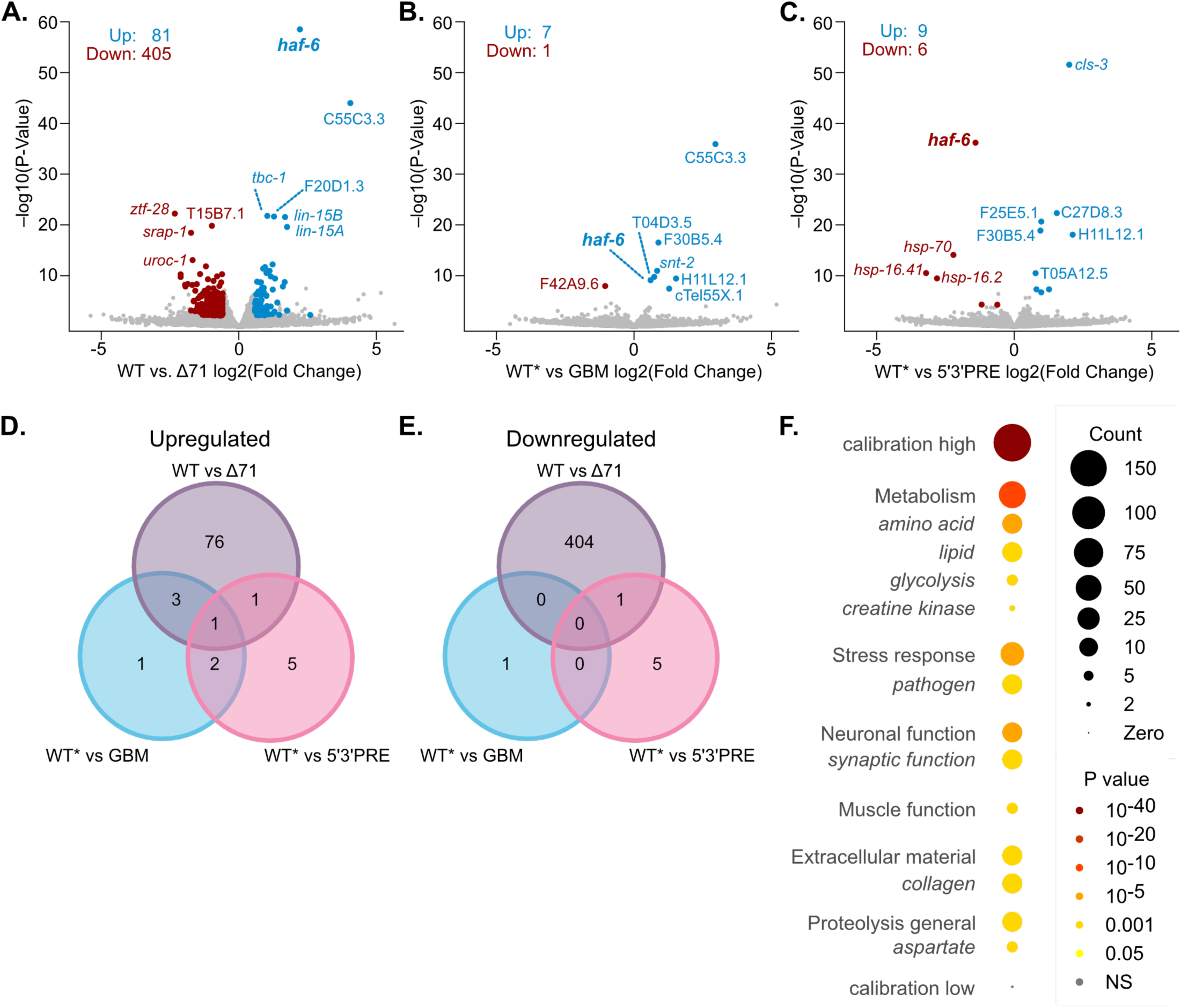
RNA-seq analysis of staged young adults including all strains from figure 4A. **A.** Volcano plot of *glp-1(WT)* vs. *glp-1(Δ71).* Gray circles indicate genes where the expression does not change or is not significant in a DESeq2 analysis comparing mutant to wild type expression, with a log_2_ fold change cut off greater than 0.585, less than -0.585, and a p_adj_ threshold of less than 0.05. Increased genes are labeled in blue, and decreased genes are labeled in red. The total number of differentially expressed genes is given. **B.** Volcano plot as in (A), except *glp-1(GBM)* gene expression is compared to *glp-1(WT*)*. **C.** Same as in (B), except *glp-1(PRE)* gene expression is compared to *glp-1(WT*)*. **D.** The overlap in the identity of differentially upregulated genes is compared between all three mutants in the Venn diagram. The circles are colored by genotype as in figure 4A. **E.** A Venn diagram as in (D), except downregulated genes are compared. **F.** Wormcat 2.0 analysis of the total regulated gene set in *glp-1(Δ71)* vs. *glp-1(WT)*. Category 1 (non-italics) and 2 (italics) changes are grouped by pathway. The size of the circle indicates the counts of genes in each category, and the color represents the P-value from the Wormcat 2.0 analysis.

The data show that the number of gene expression changes correlates with the magnitude of the impact on reproductive fecundity, and that most gene expression changes observed in *glp-1(Δ71)* are unique to that mutant. The abundance of *glp-1* mRNA is unchanged in all three mutants, suggesting that *glp-1* mRNA abundance is not impacted by either binding site mutation or by the larger deletion.

Focusing on the *glp-1(Δ71)* mutant, we used WormCat 2.0 to determine which gene sets are perturbed in this mutant compared to control (**Fig. 7F, Supplemental Data Set 2**) [55]. The results show that metabolism and stress response genes are overrepresented in the total regulated gene set (both up and downregulated genes, p_adj_ = 2.7e-13 (metabolism); p_adj_ = 3.3e-7 (stress response)). Specifically impacted metabolic pathways include amino acid metabolism, lipid metabolism, creatine kinase, and glycolysis. We also observe changes in somatic genes involved in neuronal and muscle function, although at a lower frequency (p_adj_ = 7.7e-8 (neuronal); p_adj_ = 1.5e-6 (muscle)). Using tissue enrichment analysis on upregulated genes, we see an enrichment for genes expressed in the germline (Q-value = 0.00014) and oocytes (Q-value = 0.012, **Supplemental Data Set 2**), consistent with ectopic mitosis in the germline [56]. Repeating the tissue enrichment analysis on downregulated genes, we see enrichment of genes expressed in several muscle lineages (MS, D, and C lineage), as well as the head mesodermal cell (MS) and the VC2 and VC3 neurons (P3, P5 lineages, **Supplemental Data Set 2**). The results suggest the possibility of posterior patterning defects in *glp-1(Δ71)* embryos, possibly caused by dysregulation of GLP-1 in the mutant. These changes may account for the observed reduced fecundity in this mutant, but more work will be necessary to understand the precise mechanism.

## DISCUSSION

The regulation of *glp-1* mRNA has served as a paradigmatic example of post-transcriptional control of maternal transcripts for over three decades. A pioneering study demonstrated that the *glp-1* 3ʹUTR is sufficient to pattern the expression of injected reporter mRNAs [4]. Subsequent work using reporter transgenes confirmed this observation [16], and specific functional elements within the 3ʹUTR were mapped using both approaches [4, 29, 32]. Here, we expanded upon these foundational studies in two ways. First, we evaluated the mechanism of regulation for two functional elements recognized by the RNA-binding proteins GLD-1 and POS-1 across the oocyte-to-embryo transition. Second, we evaluated three alleles corresponding to previously identified functional regions of the 3ʹUTR, defining their importance to reproductive fecundity.

Our work suggests that endogenous *glp-1* mRNA undergoes a shortening of the polyA tail as transcripts move from the adult germline to embryos. Loss of the GLD-1 or POS-1 binding motifs (GBM and 5ʹ3ʹPRE) prevents this shortening, consistent with our prior work that demonstrated an increase in reporter expression in the posterior of the embryo in reporter transgenes [32]. The increase in fluorescence observed in both mutants depends upon the cytoplasmic polyA polymerase GLD-2, suggesting that the mechanism of repression in embryos includes prevention of A-tail lengthening in embryos by this enzyme. Consistent with that model, the average polyA tail length of GBM and 5ʹ3ʹPRE mutant reporters is longer than WT embryos. Intriguingly, knock down of a second cytoplasmic polyA polymerase, GLD-4, causes the opposite effect. Our findings suggest that GLD-4 plays a repressive role in embryos, working in opposition to GLD-2, but the exact mechanism remains unknown. Knock down of GLD-4 had no impact on the 5ʹ3ʹPRE, suggesting the possibility that GLD-4-dependent repression acts through the POS-1 binding site, either directly through POS-1 or via an indirect mechanism. Our work also shows that none of the deadenylase enzymes are required for *glp-1* regulation, at least in oocytes, although this observation is limited by the potential for functional overlap between multiple deadenylase enzymes. Lastly, we observe that the atypical eIF4E homology *ife-3* represses all three GLP-1 reporters in oocytes, consistent with models indicating a translational control mechanism in the germline [57]. Together, the data shows that multiple mechanisms contribute to *glp-1* repression, and the predominant mechanism changes from oocytes to embryos.

Our work also shows that precise mutations in the GLD-1 and POS-1 binding sites do not cause strong reproductive or molecular phenotypes when engineered into the endogenous *glp-1* locus. There are several potential explanations for this apparent dichotomy. First, there could be functional redundancy between the binding sites, although the reporter transgene data support a model where binding of both proteins is needed for full repression. Second, additional mechanisms that act on GLP-1 protein, but not on a GFP reporter, may buffer modest changes in GLP-1 expression. Lastly, it is possible that additional binding sites in the coding sequence of *glp-1* mRNA could compensate for loss of the binding sites in the 3ʹUTR. The finding that a larger deletion that removes all three motifs confers a much stronger reproductive phenotype supports the first hypothesis.

How does dysregulation of *glp-1* lead to embryonic lethality? Our RNA sequencing data provide a few clues. The tissue enrichment analysis reveals that genes from the C, D, and MS lineage muscle tissues are preferentially downregulated in *glp-Δ71* mutant embryos. All three lineages are posterior in origin, deriving from cells that normally do not express GLP-1 protein. It is possible that loss of posterior lineage cell specification could account for the changes we observe in these tissues. Similarly, amino acid metabolism and creatine kinase activity is associated with muscle development and activity. The reduction we observe in genes expressed in these pathways could be explained by reduced development of muscle tissue in embryos. Consistent with this hypothesis, most *glp-1(Δ71)* mutant embryos fail before gastrulation, consistent with cell fate specification defects. We note that the terminal phenotype differs both in the timing of embryonic arrest and in the appearance of the terminal embryos from a *glp-1* null embryos, which fail during morphogenesis with anterior pharyngeal tissue defects [22].

Specific metabolism genes that are downregulated in our dataset include *pah-1*, *ceeh-2* and Y51H4A.7. The *pah-1* gene is involved in anabolism [58]. The *ceeh-2* gene is an epoxide hydrolase that functions in lipid signaling [59]. The human UROC1 homolog Y51H4A.7 is thought to operate in histidine degradation [60]. We also note that stress response genes appear to be downregulated in the mutant. Among the downregulated stress response genes are *cdr-2*, *clec-48*, and F35E12.6, C49C8.5, and ZK6.11.1. The *cdr-2* gene is involved in the response to cadmium [61]; whereas F35E12.6, C49C8.5, ZK6.11.1, and *clec-48* genes have reported roles in the innate immune response [62].

Conversely, the upregulated gene set is limited to germline and/or oocyte expressed genes that mostly have unassigned functions. The most upregulated gene is C55C3.3, a membrane-associated protein of unknown function that appears to be broadly conserved in nematodes in a BLASTP analysis [63]. This gene encodes a prion like (Q/N) domain with 41 glutamines and 27 asparagines. The protein also harbors a serine-proline rich region, encoding 57 serines and 58 prolines. This transcript is also upregulated in the *glp-1(GBM)* mutant, but not in the *glp-1(5ʹ3ʹPRE)* mutant. Knock down of the C55C3.3 in a *rrf-3* sensitized genome-wide RNAi screen causes embryonic lethality [64]. However, a loss of function allele does not appear to have this phenotype. Additional upregulated genes include *haf-6*, *lin-15A*, *lin-15B* and *tbc-1*. *tbc-1* was reported to have functions in innate immunity [65]; *haf-6* is a member of ATP-binding cassette transporters and found to be necessary for RNA interference [66]. *lin-15A* and *lin-15B* have functions in vulval development [67]. More work is needed to discern which of these changes, if any, contribute to the phenotypes observed in *glp-1(Δ71)*.

We were initially surprised that loss of the GLD-1 or POS-1 binding motifs did not cause stronger phenotypes. In fact, we expected to have difficulty recovering these mutants because we anticipated strong upregulation of GLP-1, which is known to cause a gain-of-function sterility phenotype where the germline forms a tumor of ectopic progenitor cells. This turned out not to be the case. Mutation of all three binding sites and the adjacent regions caused an increase in the amount of mitosis and a moderate expansion of the mitotic zone, but apparently the extent of dysregulation observed was not enough to cause germline progenitor cell tumor formation. Perhaps a larger deletion of the 3ʹ UTR could cause a stronger phenotype. We attempted to make a full 3ʹUTR deletion of the *glp-1* gene, but we were unable to recover this mutant, although technical hurdles associated with non-allelic homologous recombination with a downstream pseudogene appears to be the root cause of this difficulty (not shown). Having said that, we note that similar 3ʹUTR deletion alleles of key cell-fate specifying maternal genes *mex-3* and *pos-1* also do not produce strong reproductive phenotypes [49, 50]. Together with the *glp-1* results reported here, our work supports an emerging model where silent maternal transcripts inherited by embryos act to buffer protein production but are not essential for fertility.

## METHODS

### C. elegans strains and CRISPR mutagenesis

All strains used in this work were cultured on NGM agar plates seeded with *E. coli* OP50 strain [68] under standard conditions unless otherwise noted. The strains used in this study are listed in **Table 1**. CRISPR/Cas9 mutagenesis was carried out following the protocol of Ghanta et al. [69]. A single crRNA and an ssODN or DNA oligonucleotide repair template unique to each mutation was employed for the *glp-1::2xOLLAS* background mutations (*spr16, spr21, spr22, and spr33*). The sequences of the crRNA, the DNA repair template, and the ssODNs are listed in **Supplemental Table 1**. The offspring of injected animals were isolated, allowed to propagate, then lysed and used as a template in PCR using primers F21: 5ʹ-CGGTCAATGCGATGTGGTAT-3ʹ and R21-2: 5ʹ-GAGCCTCGAGAATGGTGAGT-3ʹ. For *spr16*, the presence of the deletion product was visualized directly on a 2% agarose gel. For point mutations, the amplification product was treated with the HindIII-HF restriction enzyme, where lack of cleavage indicated a successful mutation (NEB, catalog number: R3104S, New Ipswich, MA). Cleavage products were resolved on a 2% agarose gel. Sanger sequencing was used to validate the mutation after candidate homozygous lines were isolated. Validated mutants were outcrossed three times with the background strain to eliminate off target and spurious background mutations for each strain respectively.

### RNAi by feeding

Double-stranded DNA sequences used to target *gld-2* or *gld-4* in RNAi experiments were amplified from synthetic DNA (GenScript, Piscataway, NJ). Amplicons were designed using the sequences from Ahringer RNAi library [64, 70]. The *gld-2* amplicon is 1194 nucleotides (primers: *gld-2*F: 5ʹ-AGTACGGCCGTTGGTATCAG-3ʹ, *gld-2*R: 5ʹ-ATGTTGCTGCTGATTGTTGC-3ʹ, **Supplemental Table 1**), while the *gld-4* amplicon is 969 nucleotides (primers: *gld-4*F: 5ʹ-TTTCGGAGCGAGAAGACACT-3ʹ, *gld-4*R: 5ʹ-GTGGCTGAGTTGTTGTCGAA-3ʹ) [70, 71]. PCR products were cloned into vector L4440, and RNAi experiments were carried out as described previously [32, 72]. Briefly, adult worms were bleached to recover embryos, which were then placed on plates with 100 μg/mL ampicillin and 1mM IPTG to induce RNAi upon hatching. Animals were cultivated at 25°C until reaching the young adult stage. After dissection, embryos were placed on 2% agarose pad and imaged under DIC optics and fluorescence optics (see below). The previously reported phenotypes were used to confirm RNAi efficacy [44]. *pos-1* RNAi was used as a positive control condition and resulted in embryonic lethality [32].

### RNAi by soaking

Total RNA was isolated from one or two 60 mm plates of N2 worms using Trizol followed by phenol-chloroform-isoamyl alcohol extraction. RT-PCR was performed using Superscript III One-Step RT-PCR kit (ThermoFisher, catalog number: 12574026, Waltham, MA) using the primers listed in **Supplemental Table 1**. Primers for each of the genes of interest targeted exon-exon junctions to reduce amplification of residual contaminating genomic DNA. Each primer includes a T7 promoter on the 5ʹ end. The amplicons vary in length from 550 to 950 base pairs. Double stranded RNA was made from the RT-PCR products using the MEGAScript T7 Transcription Kit (ThermoFisher, catalog number: AM1333, Waltham, MA). Soaking solution was made following the procedure of Ahringer with a final concentration of 1 µg/µl of double-stranded RNA [70]. Synchronized L1 animals were incubated in this solution at 25°C for 24 hours. After transferring to NGM agar plates seeded with *E. coli* OP50, they were cultured at 25°C until the young adult stage. Embryos were harvested by dissection for imaging under DIC and fluorescence optics (see below). The previously described phenotypes for each RNAi condition were used to confirm knockdown [73].

### Brood size and hatch rate assays

Following synchronization, worms from each tested genotype were cultivated at room temperature before being isolated onto NGM plates with *E. coli* OP50. Every day, each animal was moved to a different plate, and the number of eggs laid and hatched progeny were counted. Three biological replicates including 30-35 worms were performed for each strain and culture condition. A one-way ANOVA was used to assess differences between the brood size and hatch rate (viable progeny / total progeny) with Bonferroni correction for multiple hypothesis testing.

### Reporter transgene fluorescence microscopy and quantification

A Zeiss Axioskop 2 plus microscope (Jena, Germany) with a 40X oil-immersion objective and a Spot camera (Sterling Heights, MI) was used to collect embryo images under DIC and fluorescence optics. For embryo imaging, the mean pixel fluorescence intensity in the nucleus was determined for each nucleus in all imaged four-cell embryos using FIJI v2.9.0 (https://imagej.net/software/fiji/). The reported value is the sum of the four nuclei in each four-cell embryo. The mean and standard deviation of this sum was calculated for multiple embryos under each treatment condition (**Fig. 2**). The data were plotted using Igor Pro9 (Wavemetrics, Portland, OR); and the Tukey method was used to identify outliers. A one-way ANOVA was performed to assess statistical significance, the p-value was adjusted using the Bonferroni correction for multiple hypothesis testing. For oocyte imaging, the background-corrected mean pixel fluorescence intensity in the nucleus of the four most proximal oocytes was determined. The mean pixel intensity of each was determined, and the reported value is the average of all four nuclei. The average and standard deviation from multiple worms was determined. The Tukey method was used to identify outliers, and the data were compared using a one-way ANOVA to evaluate the significance using the Bonferroni correction for multiple hypothesis testing.

### Immunofluorescence imaging and quantitation

After slides were prepared with poly-L-lysine treatment, gonads were dissected in Happy Buffer (25 mM HEPES pH7.0, 50 mM NaCl, 5 mM KCl, 2 mM MgCl_2_ and 2mM EDTA) containing 0.2 mM Levamisole. The worms were fixed with the fixation buffer (2X buffer: 25 mM HEPES pH7.4, 49 mM NaCl, 5 mM KCl, 2 mM MgCl_2_, 1mM EGTA and 5% paraformaldehyde and incubated for 30 minutes. The slides were washed with 0.2M Tris pH 7.5 twice before being incubated in Permeabilization Buffer [PBST (0.1%), Triton (0.1%)] for 15 minutes and then submerged into PBST (0.1%). After incubating with primary antibody (rat anti-OLLAS (Novus #NBP1-06713) 1:500; monoclonal mouse Phospho-Histone H3 (Ser10) (6G3) (Cell Signaling

9706S) 1:200) overnight at 4°C, PBST (0.1%) was used twice for 5 minutes to wash the slides, which was followed by the secondary antibody (AlexaFluor 488 donkey anti-rat (Invitrogen #A21208, headquarters: Carlsbad, CA) 1:500 dilution; Rhodamine Red™-X (RRX) AffiniPure Goat Anti-Mouse IgG (H+L) (Jackson ImmunoResearch 115-295-003, West Grove, PA) 1:200 dilution) treatment for 2 hours at room temperature. PBST (0.1%) was used twice to rinse the slides, then Vectashield Antifade Medium (Vector Labs, Newark, CA) containing DAPI was used to mount the gonads on the slide, covered with a coverslip, sealed with clear nail polish and allowed to dry for half an hour. Fluorescent microscopy was used to image the slides.

Images were collected on a Zeiss AxioObserver 7 inverted imaging system (model number:431007-9904-000, serial number:3858002009, Oberkochen, Germany) with 20X air and/or 40X oil objectives under DIC and fluorescence optics. Z-stacks of 30 images spanning the width of the gonads were collected using the green channel for OLLAS, the red channel for PH-3, and the blue channel for DAPI, and DIC for light microscopy. The images were quantified using Fiji version 2.14.0. The lengths of the mitotic zone and the length of the OLLAS-positive region were measured using the segmented line tool. The ollas immunofluorescence intensity was determined using the circle tool with radius of 30 pixels and normalized to an equivalent circle outside of the gonad to correct for background. The statistical significance was determined using a one-way ANOVA with Bonferroni correction for multiple hypothesis testing.

### Analysis of polyA-test sequencing data (PAT-Seq)

Raw sequencing reads from published polyA-test sequencing data corresponding to NIH Bioproject PRJNA248698 were downloaded from the NIH Sequence Read Archive [41]. Reads that correspond to *glp-1* isoforms were recovered from the raw data using the fuzzy pattern search algorithm ugrep v7.4.1 (https://github.com/Genivia/ugrep) built into a custom script (**Supplementary Data Set 3**). The pattern used for the search consists of 1) a 20 nucleotide long 3ʹUTR isoform specific search term near the polyA processing site (search1 = 5ʹ-CTCGGTTCATTTTAAATATG-3ʹ; search2 = 5ʹ-CATTTTTCTTATTCTAGACT-3ʹ; search3 = TCCTTTTCTTTATAACTTGT-3ʹ; search4 = 5ʹ-CATTTAATGAATTGTAATTC-3ʹ; search5 = 5ʹ-AGATTAAGAGTATAAGCTTT-3ʹ; search6 = 5ʹ-TTCGAATTATTGAAGCTCAA-3ʹ), 2) a variable length gap ranging from 0 to 10 nucleotides, 3) a stretch of continuous adenosines at least five nucleotides long, and 4) eight nucleotides of the tag sequence used to clone the polyA tails during library construction (GATCGGAA) [74]. The advantage of ugrep compared to similar grep-based search algorithms is that it can search for near matches. We ran our script with 0-3 possible mismatches compared to our search expression and found that 2 or 3 mismatches permitted the recovery of reads that do not correspond to *glp-1*, while 0 or 1 errors produced only matches to *glp-1* and comparable results (**Fig. 2, Supplemental Data Set 3**). The length of the polyA tail was counted using a second iteration of ugrep to recover the polyA tail with the 3ʹ tag, and awk v.20200816 was used to count the length of the polyA tail. The full script with list of commands and the output is available in **Supplemental Data Set 3**. The number of isoform-specific reads was divided by the total number of *glp-1* reads to determine the fraction of polyA tails associated with a particular UTR isoform. The mean, median, and standard deviation of each distribution was calculated. A non-parametric Komolgorov-Smirnov (KS) test was used to evaluate apparent differences between distributions, and p-values were adjusted for multiple hypothesis testing using the Bonferroni correction.

### Reporter transgene PolyA tail length measurements

Young adult animals were washed with M9 several times, centrifuged, and resuspended in Trizol (Invitrogen, catalog number: 15596018, Carlsbad, CA), frozen and stored at -80°C. To isolate embryos, young adult animals were bleached with hypochlorite solution, washed with M9, centrifuged, resuspended in Trizol, frozen and stored at -80°C. RNA was isolated by using phenolchloroform extraction method. PolyA tail length measurements were conducted as published with a few changes [45]. Superscript III One-Step RT-PCR kit (ThermoFisher, catalog number: 12574026, Waltham, MA) was utilized after the guanosine and inosine extension step [45]. The PCR fragments were run on agarose gel. Primers used in this assay: C10T2 (sequence: 5ʹ-CCCCCCCCCCTT-3ʹ) to make cDNA, M13RC0T2 (sequence: 5ʹ-CAGGAAACAGC-

TATGACCCCCCCCCCCTT-3ʹ) [45] to amplify the tail, forward primer (sequence: 5ʹ-GCGATGGCCCTGTCCTTTTA-3ʹ) to amplify both transcripts, reverse primer for the gene (sequence: 5ʹ-GGGACCAGTGTCTGGGTGATGT-3ʹ) to amplify only gene specific transcripts. TOPO Cloning was performed using StrataClone PCR Cloning Kit manual (Agilent, catalog number: 240205), which was followed by Sanger sequencing by Elim Biopharm (Hayward, CA) or Eton Bioscience (San Diego, CA). The polyA tail length for each isolate was counted directly from the sequencing results and plotted as a function of genotype. The mean, median, and standard deviation of each genotype and age group was calculated, and differences in the distributions were assessed by a non-parametric KS test with Bonferroni correction for multiple hypothesis testing.

### Terminal embryo phenotyping

Embryos from *glp-1(WT)*, *glp-1(Δ71)*, and *glp-1(ts)* strains were recovered by washing them off NGM agar plates with M9 followed by bleaching in hypochlorite solution (20% bleach, 0.25M NaOH final concentration). After washing with extensively with M9, one drop of embryos in M9 solution was placed on a 2% agarose pad on a microscope slide, covered with a cover slip, and imaged under DIC optics using a Zeiss Axiobserver 7 imaging system described above with a 63X oil immersion objective. The embryos were assessed as a function of developmental stage and defined as “normal” or “abnormal” based upon their appearance. The cross-tabulation Chisquare method was used to assess statistical significance, and p-values were adjusted for multiple hypothesis testing using the Bonferroni correction for multiple hypothesis testing.

### RNA-seq library preparation

*glp-1::2xOLLAS*, *glp-1(spr16), glp-1(spr21)*, *glp-1(spr22)* and *glp-1(spr33)* strains were synchronized by bleaching followed by hatching in M9 and arrest at the L1 stage. L1 animals were cultivated in an incubator at 20°C on NGM agar seeded with *E. coli* OP50 until they reached the young adult stage. Embryos were allowed to hatch overnight, leading to arrest at L1 stage. L1 animals were plated on NGM agar with *E. coli* OP50 and cultured in an incubator at 20°C until they reached the young adult stage. The worms were collected for RNA isolation by washing them off with M9 several times, and any extra M9 was removed. After addition of 1 mL Trizol, they were frozen at -80 °C. Phenol-chloroform extraction was used to isolate the RNA and depletion of ribosomal RNA was carried out following the procedure of Duan et al. [75]. Sample concentrations were determined using a Qubit fluorometer (ThermoFisher, Waltham, MA) and the RNA quality was assessed using a Bioanalyzer (Agilent, Santa Clara, CA). The NEBNext Ultra II RNA Library Prep kit (NEB catalog number: E7775S, New Ipswich, MA) was used to prepare the RNA-seq libraries. The libraries were indexed using NEBNext Multiplex Oligos for Illumina (Dual Index Primer Sets 1, 2, 3, and 4) (catalog numbers: E7600S, E7780S, E7710S, and E7730S). An Illumina NextSeq 500 was used to sequence the final pooled libraries. Three biological replicates were gathered for *glp-1(spr21)*, *glp-1(spr22)* and *glp-1(spr33)*, and two biological replicates for *glp-1::2xOLLAS*, *glp-1(spr16)*.

The OneStopRNAseq v1.0.0 pipeline was utilized for RNA-seq analysis, and the reads were aligned using the reference genome WBcel235 [53]. The differential gene expression was examined using the DESeq2_1.28.1 tool [54]. Cutoffs for differential expression were an absolute log2 fold change of greater than 0.585 and an FDR was less than 0.05. WormCat was used to evaluate gene function categories in differentially expressed genes [76].

## Supporting information

Supplemental Data Set 2

Supplemental Data Set 3

Supplemental Data Set 1

## Data availability

All numerical data and statistical analyses presented in this report are available in **Supplementary Data Set 1**. The output of the OneStopRNASeq pipeline, the Wormcat analysis, and the tissue enrichment analysis described in **Figure 7** are included as **Supplementary Data Set 2.** The custom scripts used to measure polyA tail length in **Figure 2**, and all output from the scripts, are included as **Supplementary Data Set 3**. Raw RNA sequencing data were uploaded to the NIH Sequence Read Archive and will be made available under Bioproject PRJNA1440940. All imaging data is available through Zenodo (DOI: 10.5281/zenodo.19183906).

## ACKNOWLEDGEMENTS

The authors thank Nick Rhind, Oliver Rando, Craig Mello, and Victor Ambros for sharing reagents and/or their advice on data analysis and interpretation. We thank Sharon Noronha and Haik Varderesian for helpful comments on the manuscript. This work was supported by NIH Grant R01GM145062 to S.P.R.

## AUTHOR CONTRIBUTIONS

PC collected all the data presented in the manuscript and made strains WRM73, WRM83, WRM84, and WRM109. PC analyzed the data with the following exceptions—SR assisted with some quantitation and statistical analyses described in Figures 2–4, and 6. All experiments were conceived by PC and SR, and both authors prepared the manuscript.

## SUPPLEMENTAL INFORMATION

**Supplemental Table 1.**
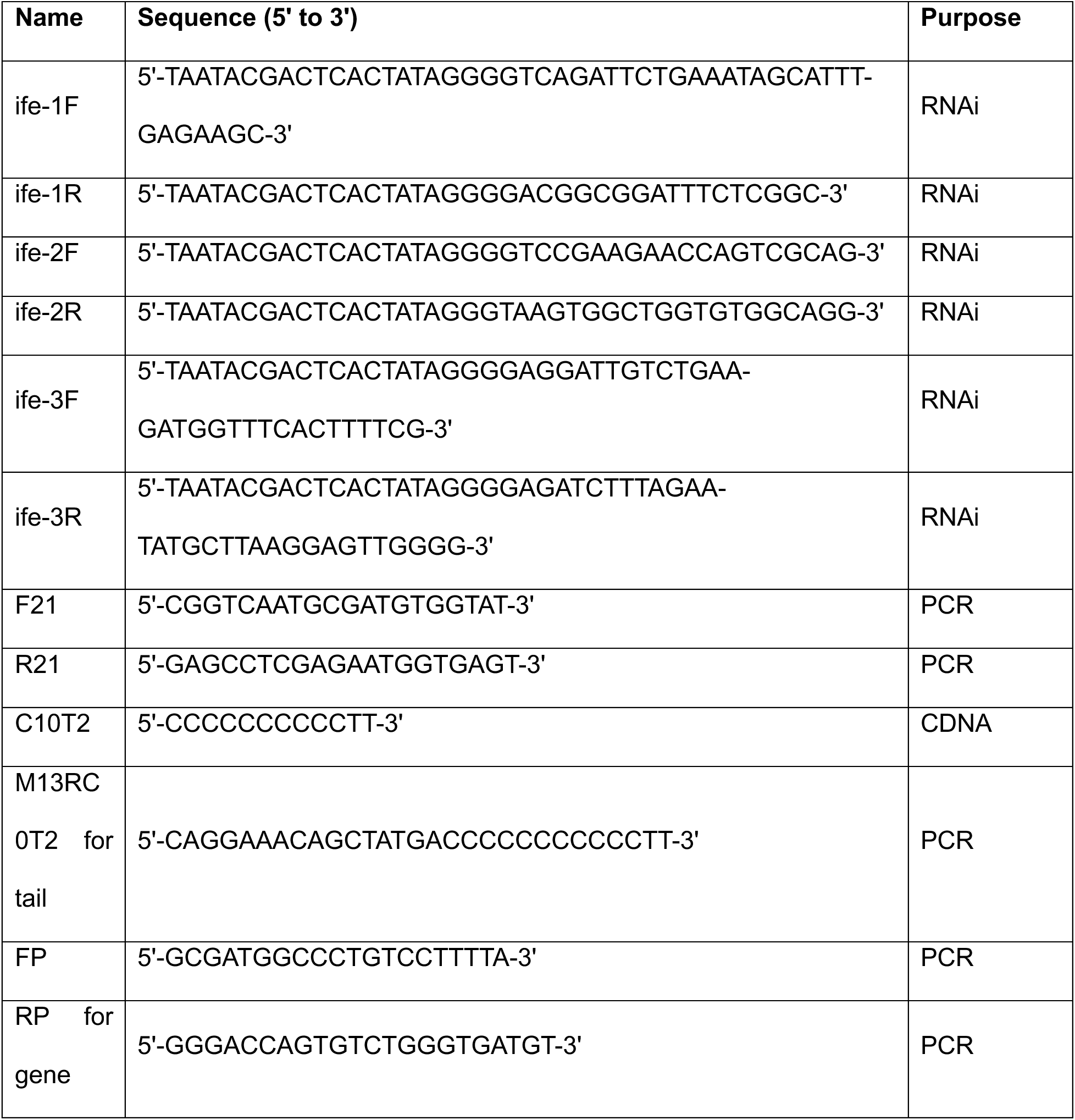
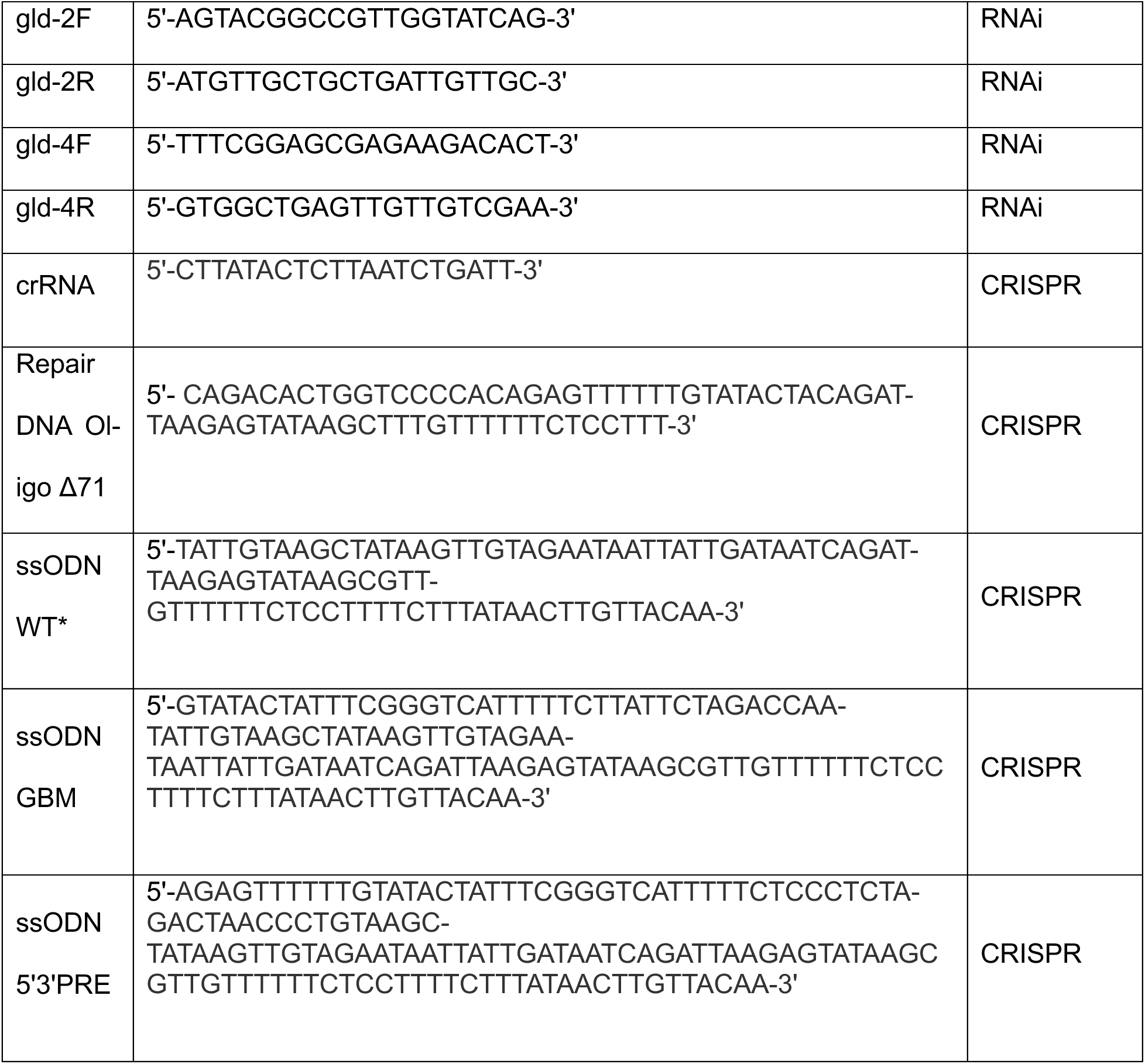
Primers and oligonucleotides used in this study.

