## Supplemental Data Set 2 for "The *glp-1* 3ʹ untranslated region regulates germline proliferation and promotes reproductive fecundity through multiple mechanisms": Enrichment Analysis.pdf

Click here for TEA results.
Click here for PEA results.
Click here for GEA results.

### Tissue Enrichment Analysis Results

| Term | Expected | Observed | Enrichment Fold Change | P value | Q value |
|---|---|---|---|---|---|
| ventral left quadrant body wall muscle WBbt:0005818 | 14 | 55 | 4 | 1.2e-18 | 8e-16 |
| body wall WBbt:0005742 | 15 | 55 | 3.7 | 3.1e-17 | 1e-14 |
| hyp12 WBbt:0004376 | 21 | 62 | 2.9 | 5.4e-14 | 1.2e-11 |
| Cpappd WBbt:0006100 | 9.3 | 34 | 3.7 | 2.9e-11 | 4.7e-09 |
| Cpapaa WBbt:0006231 | 9.3 | 34 | 3.6 | 3.2e-11 | 4.7e-09 |
| Cpapap WBbt:0006374 | 9.3 | 34 | 3.6 | 3.2e-11 | 4.7e-09 |
| head mesodermal cell WBbt:0004697 | 1.1e+02 | 180 | 1.7 | 3.4e-11 | 4.7e-09 |
| Caappd WBbt:0005991 | 9.4 | 34 | 3.6 | 3.5e-11 | 4.7e-09 |
| Cpaaap WBbt:0006357 | 9.5 | 34 | 3.6 | 5e-11 | 4.7e-09 |
| Caaapa WBbt:0006502 | 9.5 | 34 | 3.6 | 5e-11 | 4.7e-09 |
| vm1 WBbt:0006917 | 11 | 38 | 3.3 | 5.1e-11 | 4.7e-09 |
| Cpaaaa WBbt:0006151 | 9.5 | 34 | 3.6 | 5.5e-11 | 4.7e-09 |
| Caaaaa WBbt:0006076 | 9.5 | 34 | 3.6 | 6e-11 | 4.7e-09 |
| Caaaap WBbt:0006319 | 9.5 | 34 | 3.6 | 6e-11 | 4.7e-09 |
| Cpaapa WBbt:0005886 | 9.5 | 34 | 3.6 | 6e-11 | 4.7e-09 |
| Cpaapp WBbt:0006399 | 9.6 | 34 | 3.6 | 6.6e-11 | 4.7e-09 |
| Caaapp WBbt:0006521 | 9.6 | 34 | 3.5 | 7.9e-11 | 4.7e-09 |
| hyp2 WBbt:0004692 | 28 | 66 | 2.3 | 1.5e-10 | 5.3e-09 |
| tail hypodermis WBbt:0006978 | 28 | 62 | 2.2 | 4.5e-09 | 1.5e-07 |
| anal depressor muscle WBbt:0004292 | 32 | 66 | 2.1 | 1.8e-08 | 5.8e-07 |
| uterine muscle WBbt:0005342 | 13 | 35 | 2.7 | 4.8e-08 | 1.5e-06 |
| VC2 WBbt:0004619 | 12 | 33 | 2.7 | 1.1e-07 | 3.2e-06 |
| VC3 WBbt:0004618 | 12 | 33 | 2.7 | 1.1e-07 | 3.2e-06 |
| rectal valve cell WBbt:0005797 | 7.4 | 24 | 3.2 | 1.5e-07 | 4e-06 |
| anal sphincter muscle WBbt:0005798 | 17 | 40 | 2.4 | 2.2e-07 | 5.8e-06 |
| vulC WBbt:0006765 | 19 | 44 | 2.3 | 2.5e-07 | 6.1e-06 |
| OL sheath cell WBbt:0008413 | 31 | 61 | 2 | 3.6e-07 | 8.5e-06 |
| sex organ WBbt:0008422 | 1.4e+02 | 199 | 1.4 | 4.3e-07 | 1e-05 |
| Y cell WBbt:0004578 | 29 | 57 | 2 | 7.5e-07 | 1.7e-05 |
| vm2 WBbt:0006918 | 12 | 30 | 2.5 | 1.7e-06 | 3.8e-05 |
| midbody WBbt:0005740 | 45 | 78 | 1.7 | 1.8e-06 | 3.8e-05 |
| VC6 WBbt:0004611 | 18 | 39 | 2.2 | 2.1e-06 | 4.2e-05 |
| B cell WBbt:0003825 | 30 | 57 | 1.9 | 2.1e-06 | 4.2e-05 |
| rectal gland cell WBbt:0005799 | 40 | 69 | 1.7 | 5.6e-06 | 0.00011 |
| dorsal uterine cell WBbt:0006782 | 18 | 38 | 2.1 | 5.7e-06 | 0.00011 |
| head muscle WBbt:0006761 | 12 | 28 | 2.3 | 1.8e-05 | 0.00031 |
| VC1 WBbt:0004621 | 20 | 40 | 2 | 2.2e-05 | 0.00039 |
| ABarppaapa WBbt:0006180 | 9.3 | 23 | 2.5 | 2.9e-05 | 0.0005 |
| ABprappppa WBbt:0006226 | 9.4 | 23 | 2.5 | 3.1e-05 | 0.00051 |
| ABarppppapa WBbt:0006433 | 9.4 | 23 | 2.5 | 3.1e-05 | 0.00051 |
| ABarpaapap WBbt:0006375 | 9.4 | 23 | 2.4 | 3.6e-05 | 0.00057 |
| ABarpaappa WBbt:0005939 | 9.5 | 23 | 2.4 | 3.8e-05 | 0.00059 |
| ABplappppa WBbt:0004671 | 9.5 | 23 | 2.4 | 4e-05 | 0.0006 |
| ABpraapppa WBbt:0006244 | 9.5 | 23 | 2.4 | 4e-05 | 0.0006 |
| ABpraapppp WBbt:0006608 | 9.5 | 23 | 2.4 | 4e-05 | 0.0006 |
| ABarpaappp WBbt:0006070 | 9.5 | 23 | 2.4 | 4.2e-05 | 0.0006 |
| intestinal muscle WBbt:0005796 | 28 | 50 | 1.8 | 6.2e-05 | 0.00086 |
| ABplaapppa WBbt:0006735 | 9.8 | 23 | 2.4 | 6.3e-05 | 0.00086 |
| ABplaapppp WBbt:0006477 | 9.8 | 23 | 2.3 | 6.6e-05 | 0.00087 |
| tail precursor cell WBbt:0008409 | 9.9 | 23 | 2.3 | 7.3e-05 | 0.00094 |
| AS2 WBbt:0003924 | 6.3 | 16 | 2.5 | 0.00022 | 0.0028 |
| AS3 WBbt:0003923 | 6.3 | 16 | 2.5 | 0.00022 | 0.0028 |
| ut2 WBbt:0006786 | 6.3 | 16 | 2.5 | 0.00022 | 0.0028 |
| W cell WBbt:0004583 | 9.4 | 21 | 2.2 | 0.00022 | 0.0028 |
| outer labial quadrant sensillum WBbt:0005502 | 39 | 61 | 1.6 | 0.00031 | 0.0036 |
| uterine seam cell WBbt:0006789 | 7.1 | 17 | 2.4 | 0.00036 | 0.0041 |
| G cell WBbt:0008599 | 9.7 | 21 | 2.2 | 0.00037 | 0.0042 |
| ut3 WBbt:0006787 | 6.2 | 15 | 2.4 | 0.00056 | 0.0063 |
| ut1 WBbt:0006785 | 6.3 | 15 | 2.4 | 0.00062 | 0.0068 |
| PVD WBbt:0006831 | 94 | 123 | 1.3 | 0.0012 | 0.013 |
| RIF WBbt:0006835 | 4.3 | 11 | 2.6 | 0.0013 | 0.014 |
| AS9 WBbt:0003907 | 5.6 | 13 | 2.3 | 0.0016 | 0.017 |
| AS10 WBbt:0003906 | 5.6 | 13 | 2.3 | 0.0016 | 0.017 |
| uv2 WBbt:0006792 | 11 | 21 | 1.9 | 0.0017 | 0.017 |
| spermathecal-uterine junction WBbt:0006756 | 12 | 23 | 1.9 | 0.0017 | 0.017 |
| intestine WBbt:0005772 | 2.2e+02 | 258 | 1.2 | 0.0018 | 0.018 |
| striated muscle WBbt:0005779 | 58 | 80 | 1.4 | 0.002 | 0.019 |
| PVW WBbt:0008438 | 13 | 23 | 1.8 | 0.0024 | 0.023 |
| VA11 WBbt:0004647 | 11 | 20 | 1.9 | 0.0032 | 0.03 |
| uv1 WBbt:0006791 | 7.3 | 15 | 2 | 0.0032 | 0.03 |
| somatic gonad WBbt:0005785 | 1.1e+02 | 135 | 1.3 | 0.0034 | 0.031 |
| hyp6 WBbt:0004679 | 20 | 32 | 1.6 | 0.004 | 0.036 |
| hyp4 WBbt:0004687 | 16 | 27 | 1.7 | 0.0042 | 0.037 |
| ILshVR WBbt:0004523 | 15 | 25 | 1.7 | 0.0043 | 0.037 |
| DB6 WBbt:0004841 | 24 | 37 | 1.5 | 0.005 | 0.043 |
| DB4 WBbt:0004843 | 24 | 37 | 1.5 | 0.0051 | 0.043 |
| DB7 WBbt:0004840 | 24 | 37 | 1.5 | 0.0051 | 0.043 |
| DB3 WBbt:0004844 | 24 | 37 | 1.5 | 0.0051 | 0.043 |
| DB5 WBbt:0004842 | 24 | 37 | 1.5 | 0.0052 | 0.043 |
| VB9 WBbt:0004627 | 13 | 22 | 1.7 | 0.0056 | 0.045 |
| VB6 WBbt:0004633 | 13 | 22 | 1.7 | 0.0056 | 0.045 |
| VB8 WBbt:0004629 | 13 | 22 | 1.7 | 0.0056 | 0.045 |
| pharyngeal epithelial cell WBbt:0005459 | 14 | 24 | 1.7 | 0.0056 | 0.045 |

| | | | | | | |
|---|---|---|---|---|---|---|
| VB7 | WBbt:0004631 | 13 | 22 | 1.7 | 0.0057 | 0.045 |
| VB4 | WBbt:0004638 | 13 | 22 | 1.7 | 0.0059 | 0.045 |
| VB5 | WBbt:0004637 | 13 | 22 | 1.7 | 0.0059 | 0.045 |
| Psub1 | WBbt:0006874 | 7.5 | 14 | 1.9 | 0.0093 | 0.069 |
| DA6 | WBbt:0004861 | 21 | 32 | 1.5 | 0.01 | 0.075 |
| DA7 | WBbt:0004859 | 21 | 32 | 1.5 | 0.01 | 0.075 |
| hyp5 | WBbt:0004685 | 20 | 30 | 1.5 | 0.01 | 0.075 |
| DA8 | WBbt:0004858 | 21 | 32 | 1.5 | 0.011 | 0.076 |
| PDE socket cell | WBbt:0008421 | 20 | 30 | 1.5 | 0.011 | 0.081 |
| AS11 | WBbt:0003905 | 12 | 20 | 1.6 | 0.014 | 0.096 |

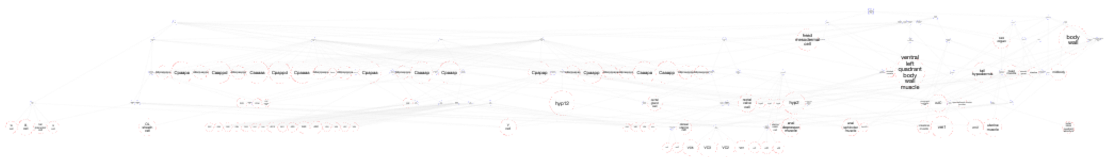

**SObA Terms - anatomy**

Save Image

Graph Depth 6

Legend :

Root

Inferred Term

Enriched Term

Inference Direction

Significance weighted

Significance unweighted

Q Value: 0.0036
WBbt:0005502 - outer labial quadrant sensillum

Return up to 15 most significant anatomy terms.

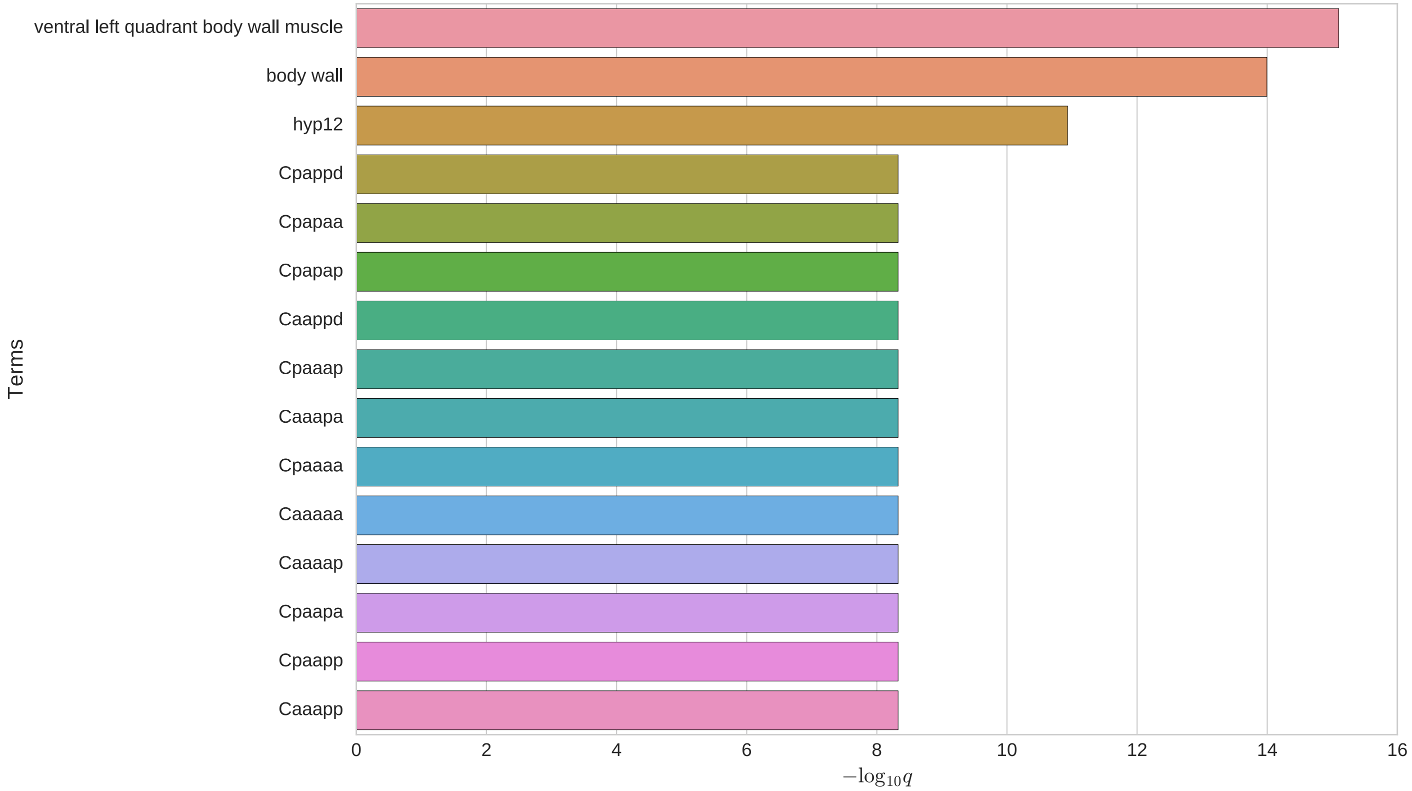


Drag graph to your desktop to save.
Download results table here.
Download observed gene table here.

Your list has 2 valid WormBase genes that have no annotated data or are excluded from testing :
WBGene00009047 - WBGene00009047
WBGene00271642 - WBGene00271642

Your list has 403 valid WormBase genes included in statistical testing :
WBGene00000007 - WBGene00000007
WBGene00000040 - WBGene00000040
WBGene00000066 - WBGene00000066
WBGene00000067 - WBGene00000067
WBGene00000136 - WBGene00000136
WBGene00000138 - WBGene00000138

perform another query

#### Phenotype Enrichment Analysis Results

| | Term | Expected | Observed | Enrichment Fold Change | P value | Q value |
|---|---|---|---|---|---|---|
| muscle system morphology variant | WBPhenotype:0000603 | 6.1 | 19 | 3.1 | 4.3e-06 | 0.0011 |
| mitochondria morphology variant | WBPhenotype:0001401 | 5.4 | 17 | 3.2 | 9.5e-06 | 0.0012 |
| body morphology variant | WBPhenotype:0000072 | 29 | 54 | 1.8 | 1.1e-05 | 0.0012 |
| body region phenotype | WBPhenotype:0002557 | 35 | 60 | 1.7 | 2.7e-05 | 0.0016 |
| sluggish | WBPhenotype:0000646 | 4.7 | 13 | 2.8 | 0.00032 | 0.016 |
| nicotine hypersensitive | WBPhenotype:0001202 | 1 | 5 | 4.8 | 0.00062 | 0.025 |
| small | WBPhenotype:0000229 | 13 | 25 | 1.9 | 0.001 | 0.036 |
| extended life span | WBPhenotype:0000061 | 16 | 28 | 1.8 | 0.0012 | 0.036 |
| paralyzed | WBPhenotype:0000644 | 3.8 | 10 | 2.7 | 0.0015 | 0.041 |
| movement variant | WBPhenotype:0001206 | 42 | 61 | 1.5 | 0.0016 | 0.041 |
| dumpy | WBPhenotype:0000583 | 8.2 | 17 | 2.1 | 0.0019 | 0.042 |
| intestinal vacuole | WBPhenotype:0001428 | 1.3 | 5 | 3.8 | 0.0022 | 0.044 |

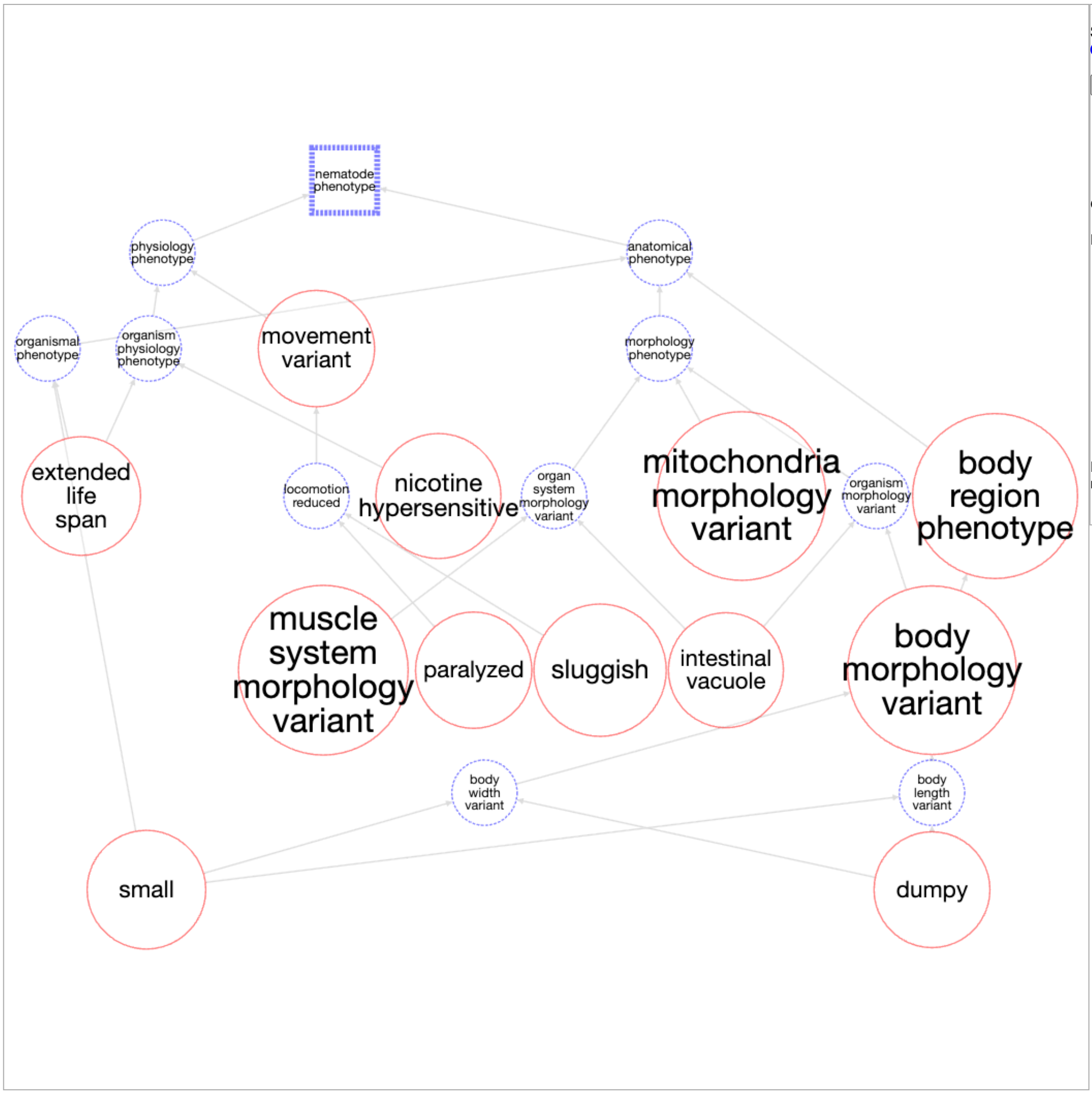


**SObA Terms - phenotype**

Save Image

Graph Depth 6

Legend :

Root

Inferred Term

Enriched Term

Inference Direction

Significance weighted

Significance unweighted

Mouseover or click node for more information.

Return up to 15 most significant phenotype terms.

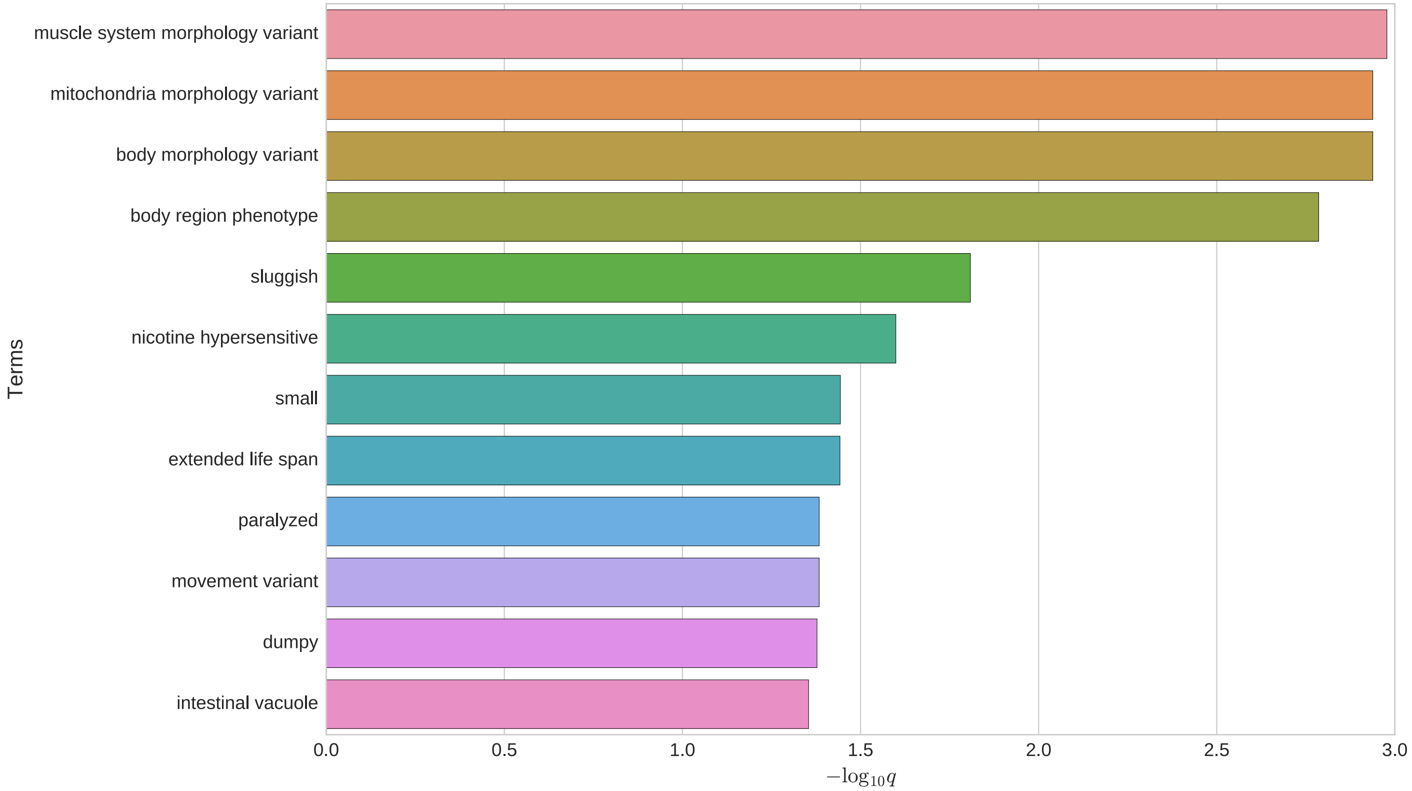


Drag graph to your desktop to save.
Download results table here.
Download observed gene table here.

Your list has 190 valid WormBase genes that have no annotated data or are excluded from testing :
WBGene00000136 - WBGene00000136
WBGene00000172 - WBGene00000172
WBGene00000175 - WBGene00000175
WBGene00000179 - WBGene00000179
WBGene00000245 - WBGene00000245
WBGene00000671 - WBGene00000671

Your list has 215 valid WormBase genes included in statistical testing :
WBGene00000007 - WBGene00000007
WBGene00000040 - WBGene00000040
WBGene00000066 - WBGene00000066
WBGene00000067 - WBGene00000067
WBGene00000138 - WBGene00000138
WBGene00000150 - WBGene00000150

perform another query

#### Gene Ontology Enrichment Analysis Results

| Term | Expected | Observed | Enrichment Fold Change | P value | Q value |
|---|---|---|---|---|---|
| extracellular region GO:0005576 | 16 | 50 | 3.1 | 8.5e-13 | 2.4e-10 |
| myofibril GO:0030016 | 2.6 | 17 | 6.6 | 1e-10 | 1.4e-08 |
| supramolecular polymer GO:0099081 | 8.2 | 30 | 3.6 | 3.9e-10 | 3.6e-08 |
| organic acid metabolic process GO:0006082 | 11 | 34 | 3.1 | 2e-09 | 1.4e-07 |
| apical part of cell GO:0045177 | 2.8 | 14 | 5 | 1.5e-07 | 8.2e-06 |
| structural constituent of cytoskeleton GO:0005200 | 0.92 | 8 | 8.7 | 2.3e-07 | 1.1e-05 |
| membrane microdomain GO:0098857 | 1.2 | 9 | 7.5 | 2.5e-07 | 1.1e-05 |
| muscle system process GO:0003012 | 2.2 | 12 | 5.4 | 3.4e-07 | 1.2e-05 |
| striated muscle dense body GO:0055120 | 2.5 | 12 | 4.9 | 1.1e-06 | 3.3e-05 |
| cuticle development GO:0042335 | 1.5 | 7 | 4.6 | 0.00013 | 0.0035 |
| collagen trimer GO:0005581 | 4.4 | 13 | 3 | 0.00014 | 0.0036 |
| response to biotic stimulus GO:0009607 | 12 | 25 | 2.1 | 0.00025 | 0.0057 |
| biological process involved in interspecies interaction between organisms GO:0044419 | 12 | 25 | 2.1 | 0.00026 | 0.0057 |
| basal part of cell GO:0045178 | 1.8 | 7 | 3.8 | 0.0005 | 0.01 |
| structural constituent of cuticle GO:0042302 | 5 | 13 | 2.6 | 0.00052 | 0.01 |
| immune system process GO:0002376 | 9.4 | 20 | 2.1 | 0.00059 | 0.01 |
| actin filament binding GO:0051015 | 2.4 | 8 | 3.4 | 0.00064 | 0.011 |
| nucleus localization GO:0051647 | 1.1 | 5 | 4.5 | 0.00075 | 0.012 |
| defense response to other organism GO:0098542 | 8.9 | 19 | 2.1 | 0.00077 | 0.012 |
| actin filament-based process GO:0030029 | 5.8 | 13 | 2.2 | 0.0022 | 0.031 |
| microbody GO:0042579 | 2.3 | 7 | 3 | 0.0023 | 0.031 |
| molting cycle GO:0042303 | 2.9 | 8 | 2.8 | 0.0025 | 0.032 |
| A band GO:0031672 | 1 | 4 | 3.8 | 0.0038 | 0.046 |
| response to bacterium GO:0009617 | 4.9 | 11 | 2.3 | 0.0038 | 0.046 |
| monocarboxylic acid biosynthetic process GO:0072330 | 1.5 | 5 | 3.3 | 0.0039 | 0.046 |
| monocarboxylic acid catabolic process GO:0072329 | 2 | 6 | 3 | 0.0041 | 0.046 |
| carbohydrate transport GO:0008643 | 1.1 | 4 | 3.7 | 0.0042 | 0.046 |
| vitamin B6 binding GO:0070279 | 1.1 | 4 | 3.6 | 0.0047 | 0.046 |
| purine nucleoside triphosphate metabolic process GO:0009144 | 1.6 | 5 | 3.1 | 0.0059 | 0.056 |

| | | | | | | |
|---|---|---|---|---|---|---|
| negative regulation of gene expression epigenetic | GO:0045814 | 1.6 | 5 | 3.1 | 0.0059 | 0.056 |
| mating behavior | GO:0007617 | 1.2 | 4 | 3.4 | 0.0063 | 0.056 |
| cellular anatomical entity morphogenesis | GO:0032989 | 1.2 | 4 | 3.3 | 0.0069 | 0.06 |
| excretion | GO:0007588 | 1.3 | 4 | 3.2 | 0.0082 | 0.069 |
| calcium ion binding | GO:0005509 | 4.1 | 9 | 2.2 | 0.0091 | 0.075 |
| unfolded protein binding | GO:0051082 | 1.3 | 4 | 3.1 | 0.0097 | 0.077 |
| muscle cell development | GO:0055001 | 1.4 | 4 | 3 | 0.011 | 0.087 |
| oxidoreductase activity acting on CH-OH group of donors | GO:0016614 | 1.9 | 5 | 2.6 | 0.012 | 0.088 |
| entry into diapause | GO:0055115 | 0.9 | 3 | 3.4 | 0.012 | 0.088 |
| iron ion binding | GO:0005506 | 3.1 | 7 | 2.2 | 0.014 | 0.1 |

**SObA Terms - go**

Save Image

Graph Depth 7

Legend :

- Root
- Inferred Term
- Enriched Term
- Inference Direction
- Inferred Alliance Slim Term
- Enriched Alliance Slim Term

- Significance weighted
- Significance unweighted

- Biological Process
- Cellular Component
- Molecular Function

Mouseover or click node for more information.

Alliance Slim terms in graph:

Return up to 15 most significant go terms.

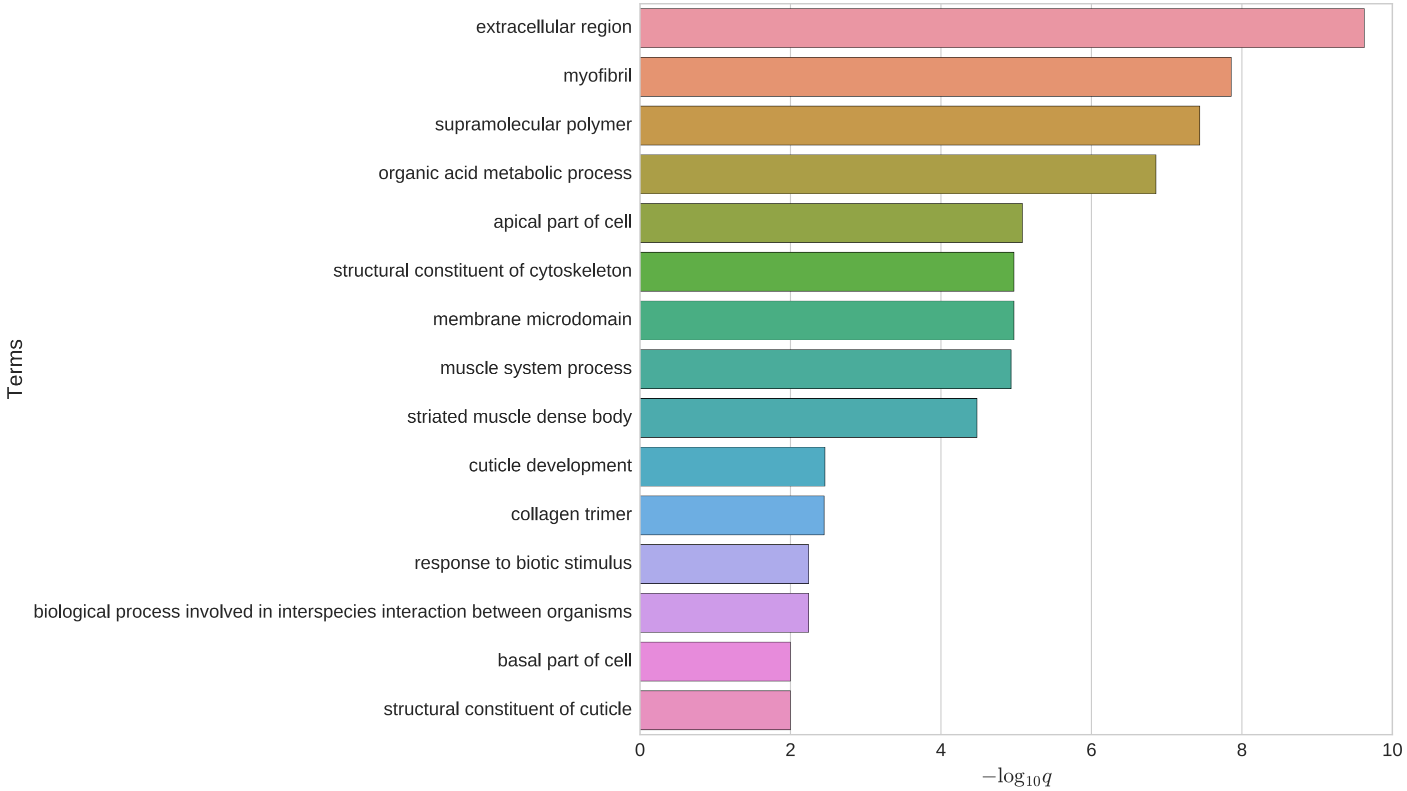


Drag graph to your desktop to save.
Download results table here.
Download observed gene table here.

Your list has 129 valid WormBase genes that have no annotated data or are excluded from testing :

WBGene00000273 - WBGene00000273
WBGene00000775 - WBGene00000775
WBGene00001000 - WBGene00001000
WBGene00001702 - WBGene00001702
WBGene00002272 - WBGene00002272
WBGene00003748 - WBGene00003748

Your list has 276 valid WormBase genes included in statistical testing :

WBGene00000007 - WBGene00000007
WBGene00000040 - WBGene00000040
WBGene00000066 - WBGene00000066
WBGene00000067 - WBGene00000067
WBGene00000136 - WBGene00000136
WBGene00000138 - WBGene00000138

perform another query
