## Supplemental Data Set 2 for "The *glp-1* 3ʹ untranslated region regulates germline proliferation and promotes reproductive fecundity through multiple mechanisms": Enrichment Analysis.pdf

Click here for TEA results.
Click here for PEA results.
Click here for GEA results.

#### Tissue Enrichment Analysis Results

| | Term | Expected | Observed | Enrichment Fold Change | P value | Q value |
|---|---|---|---|---|---|---|
| germ line | WBbt:0005784 | 30 | 61 | 2 | 2.1e-07 | 0.00014 |
| oocyte | WBbt:0006797 | 9.9 | 24 | 2.4 | 3.8e-05 | 0.012 |

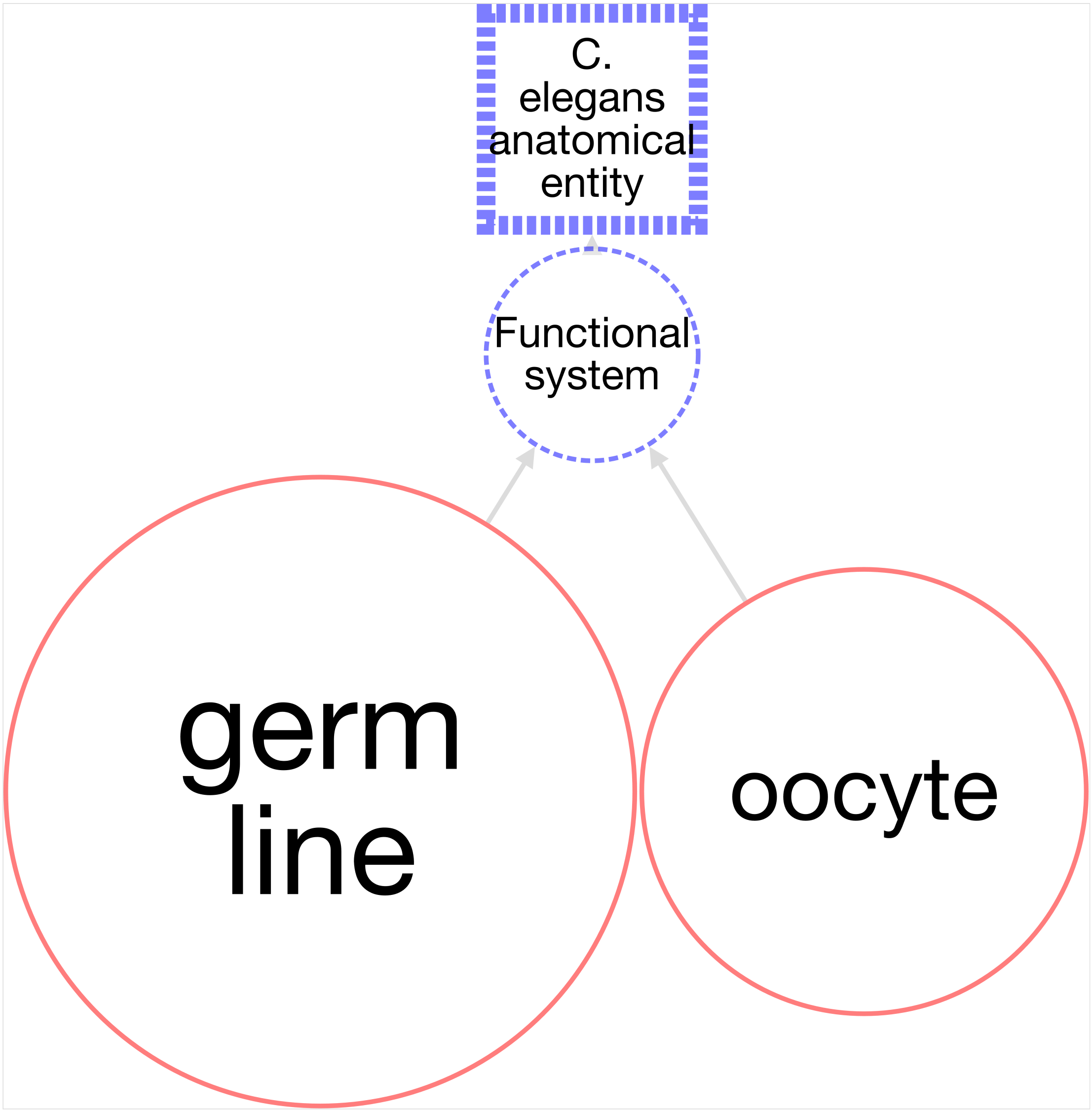


**SObA Terms - anatomy**

go back

Graph Depth 3

Legend :
- Root
- Inferred Term
- Enriched Term
- Inference Direction

drag image to desktop, or right-click and save image as

Return up to 15 most significant anatomy terms.

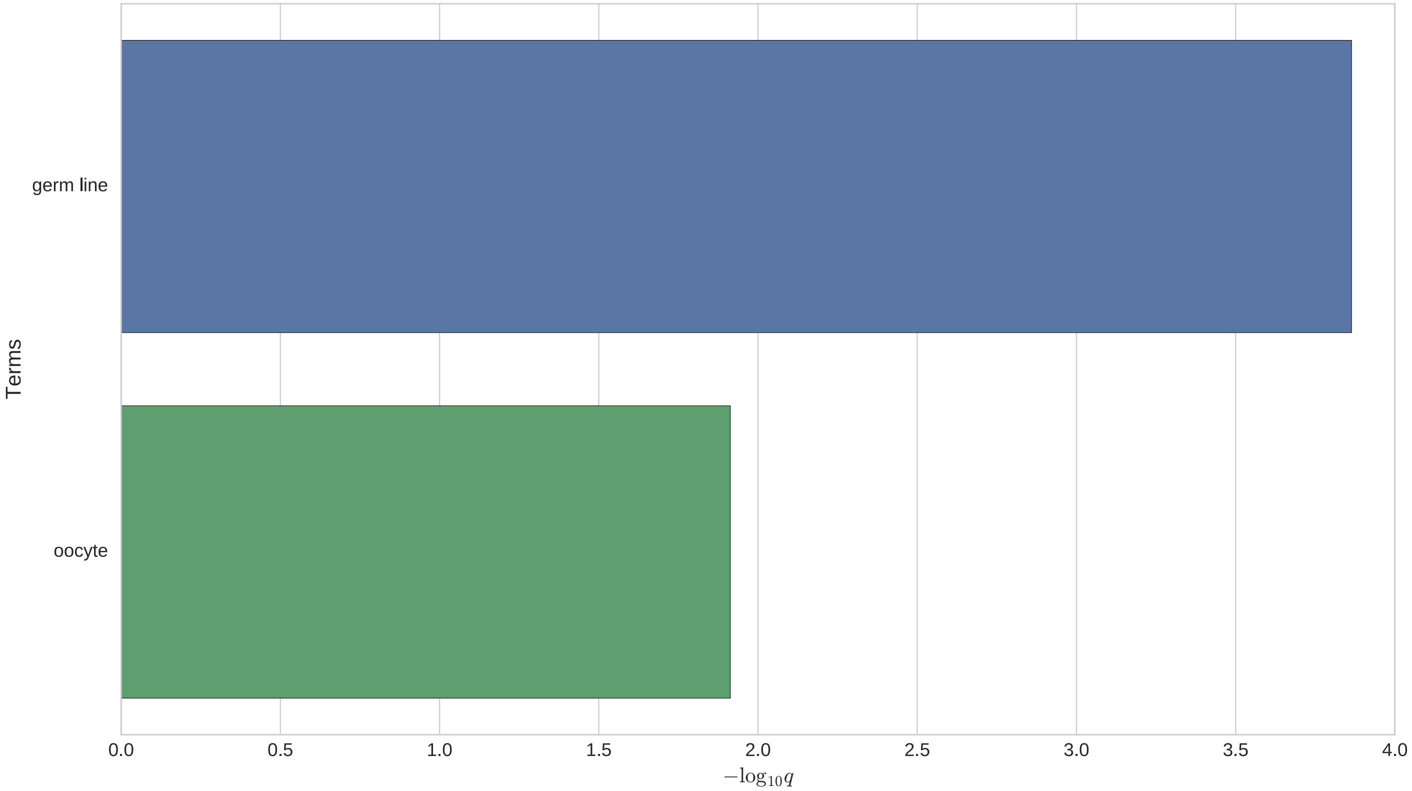


Drag graph to your desktop to save.
Download results table here.
Download observed gene table here.

Your list has 3 valid WormBase genes that have no annotated data or are excluded from testing :
WBGene00008349 - WBGene00008349
WBGene00219822 - WBGene00219822
WBGene00305050 - WBGene00305050

Your list has 78 valid WormBase genes included in statistical testing :
WBGene00000497 - WBGene00000497
WBGene00000591 - WBGene00000591
WBGene00001046 - WBGene00001046
WBGene00001130 - WBGene00001130
WBGene00001335 - WBGene00001335
WBGene00001816 - WBGene00001816

perform another query

### Phenotype Enrichment Analysis Results

| | Term | Expected | Observed | Enrichment Fold Change | P value | Q value |
|---|---|---|---|---|---|---|
| egg laying defective | WBPhenotype:0000006 | 1.3 | 6 | 4.6 | 0.00038 | 0.094 |

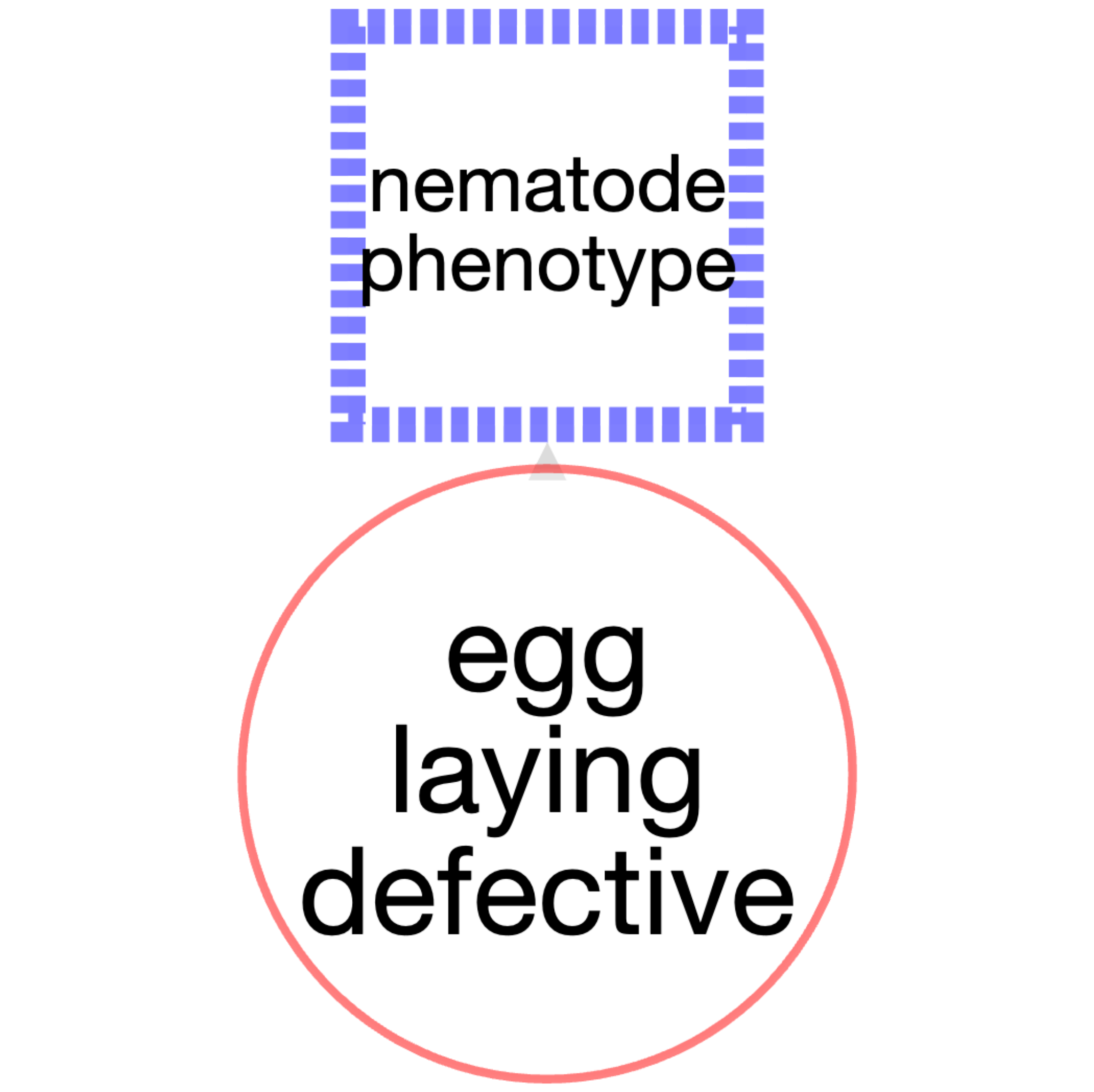


**SObA Terms - phenotype**

Save Image

Graph Depth 2

Legend :

Root

Inferred Term

Enriched Term

Inference Direction

Significance weighted

Significance unweighted

Mouseover or click node for more information.

Return up to 15 most significant phenotype terms.

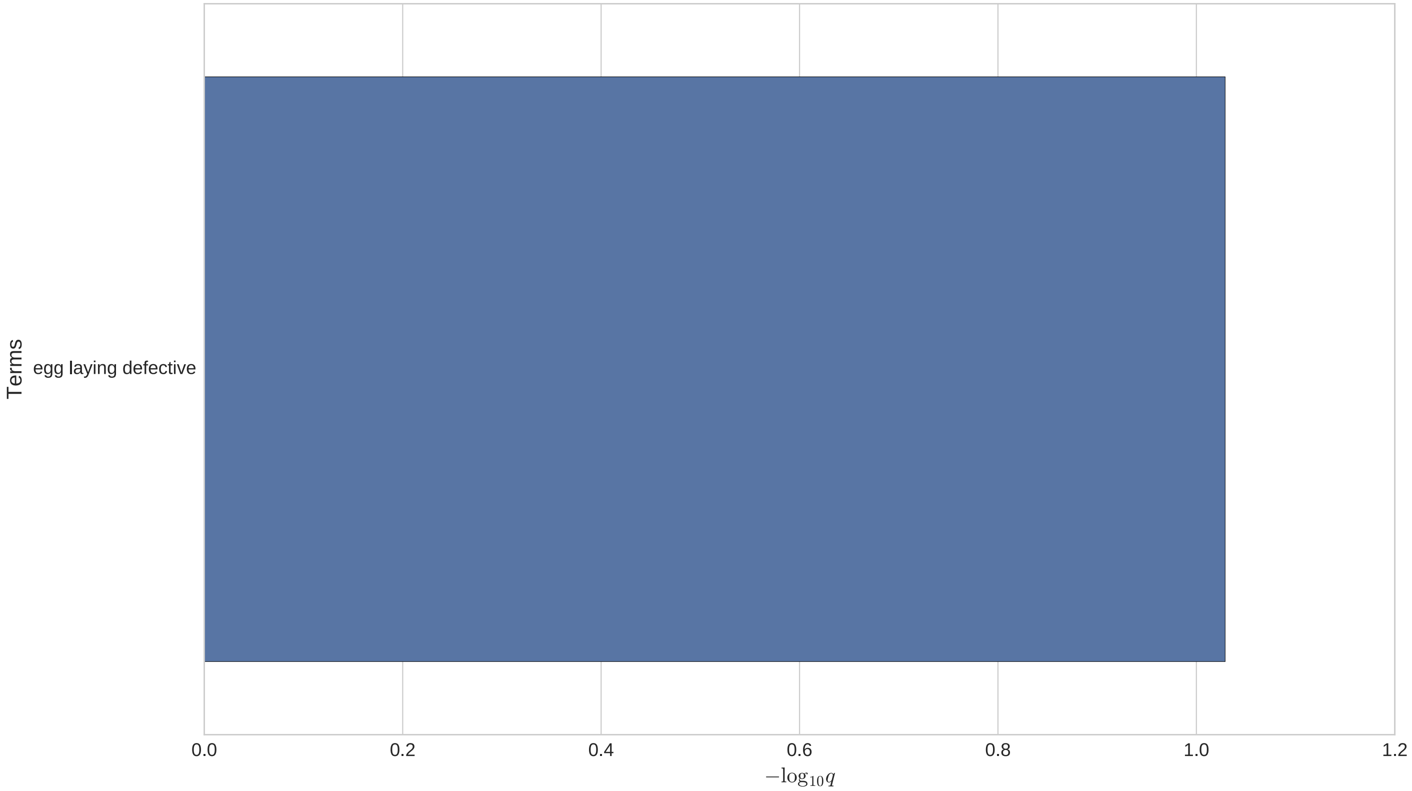


Drag graph to your desktop to save.
Download results table here.
Download observed gene table here.

Your list has 43 valid WormBase genes that have no annotated data or are excluded from testing :
WBGene00001046 - WBGene00001046
WBGene00003638 - WBGene00003638
WBGene00006404 - WBGene00006404
WBGene00007059 - WBGene00007059
WBGene00007136 - WBGene00007136
WBGene00007165 - WBGene00007165

Your list has 38 valid WormBase genes included in statistical testing :
WBGene00000497 - WBGene00000497
WBGene00000591 - WBGene00000591
WBGene00001130 - WBGene00001130
WBGene00001335 - WBGene00001335
WBGene00001816 - WBGene00001816
WBGene00001910 - WBGene00001910

perform another query

#### Gene Ontology Enrichment Analysis Results

No significantly enriched terms have been found.

Your list has 42 valid WormBase genes that have no annotated data or are excluded from testing :
WBGene00001335 - WBGene00001335
WBGene00001910 - WBGene00001910
WBGene00007124 - WBGene00007124
WBGene00007136 - WBGene00007136
WBGene00007165 - WBGene00007165
WBGene00007766 - WBGene00007766

Your list has 39 valid WormBase genes included in statistical testing :
WBGene00000497 - WBGene00000497
WBGene00000591 - WBGene00000591
WBGene00001046 - WBGene00001046
WBGene00001130 - WBGene00001130
WBGene00001816 - WBGene00001816
WBGene00003638 - WBGene00003638

perform another query
