## Supplementary figures and images for "The *glp-1* 3ʹ untranslated region regulates germline proliferation and promotes reproductive fecundity through multiple mechanisms"

### download.png

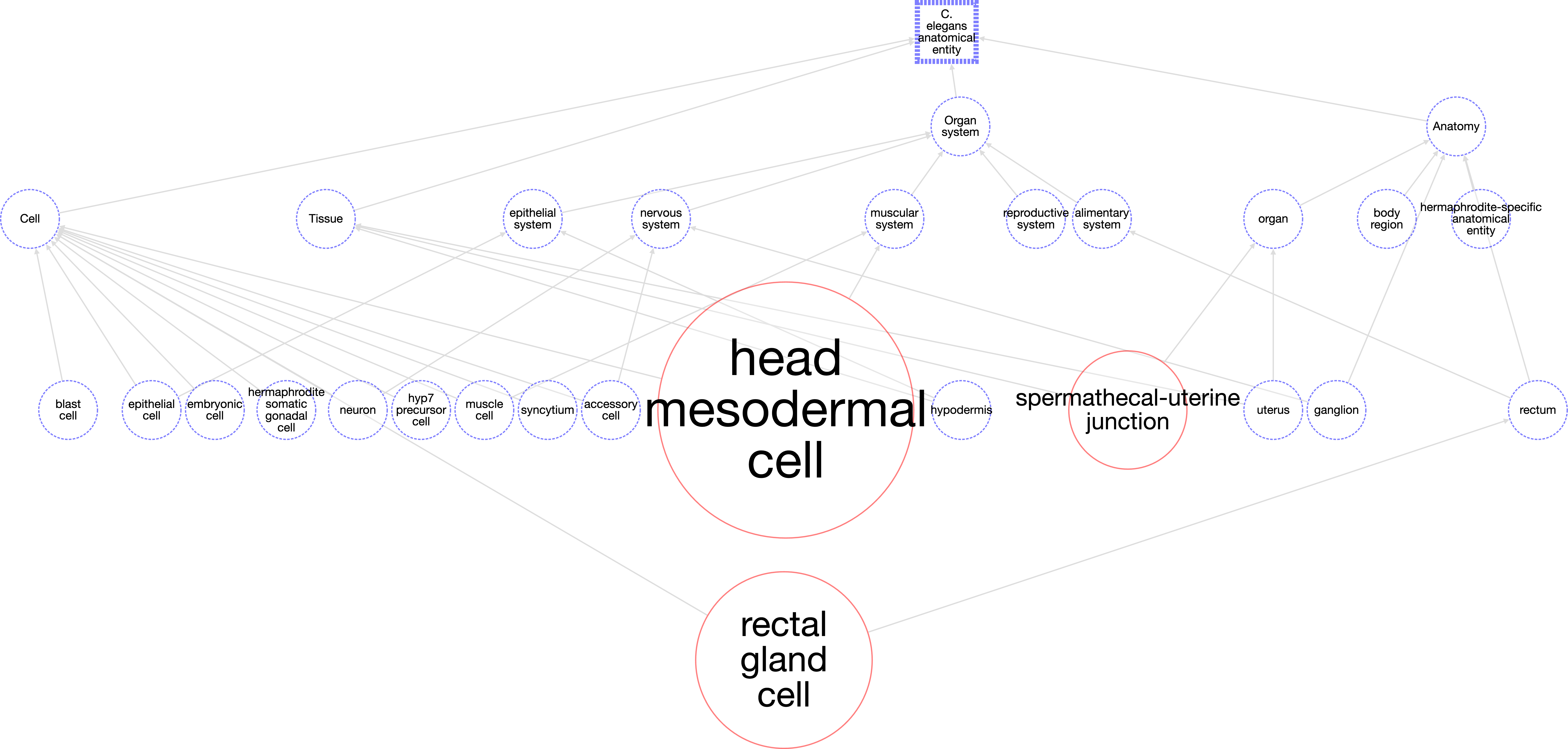

### SObA analysis.png

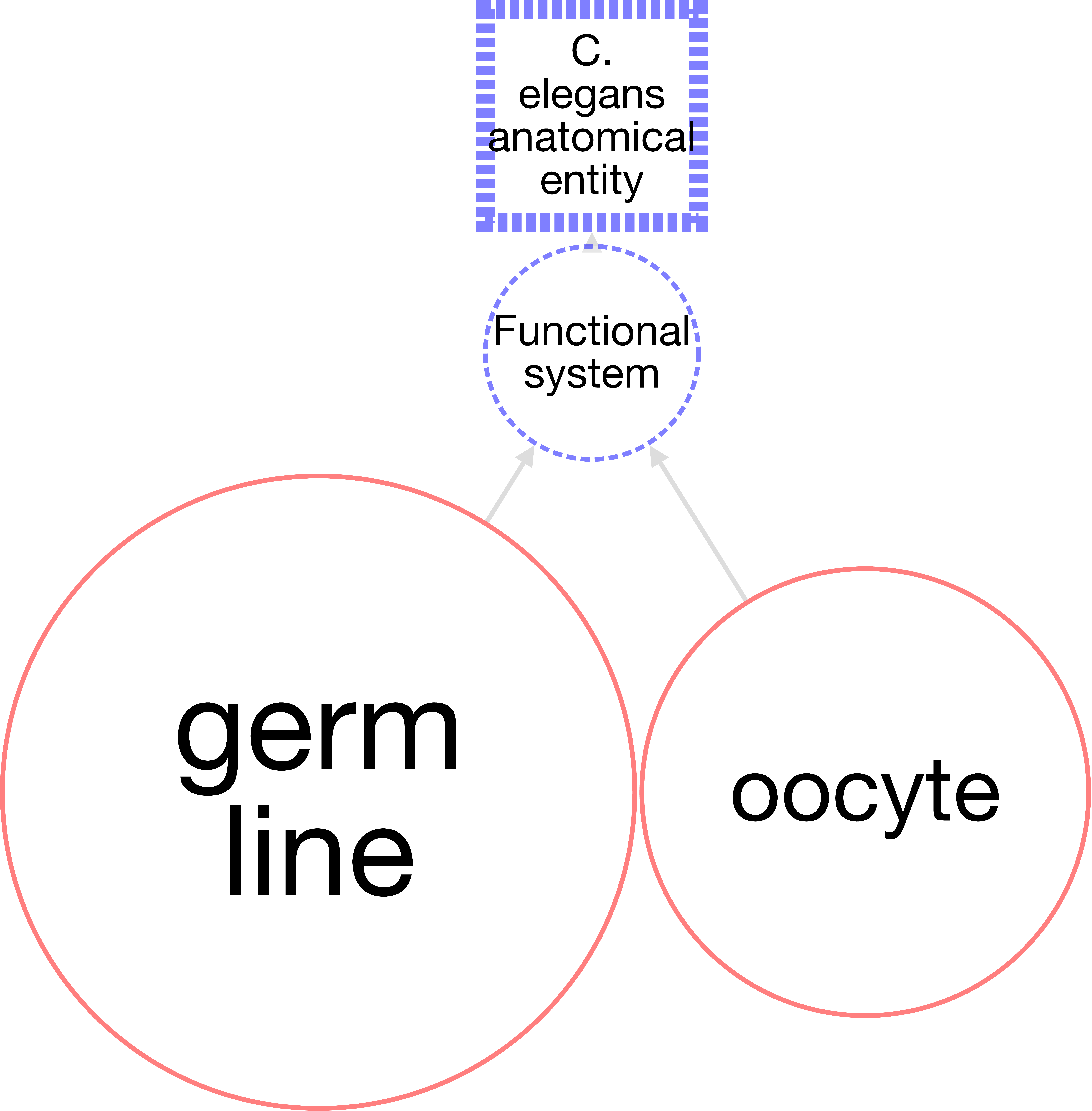
