## Supplemental Data Set 3 for "The *glp-1* 3ʹ untranslated region regulates germline proliferation and promotes reproductive fecundity through multiple mechanisms": PAtest_script.sh

```
#! /bin/bash

#Dependencies
#This script assumes that you have a working copy of ugrep installed. https://github.com/Genivia/ugrep
#This script also assumes you have two replicates of single-ended sequencing reads. Lastly, this script was designed to identify gene specific reads within bulk PolyA Test sequencing data, and incorporates sequence tags from the Harrison et al. manuscript in the search
#Harrison PF, Powell DR, Clancy JL, Preiss T, Boag PR, Traven A, Seemann T, Beilharz TH. PAT-seq: a method to study the integration of 3'-UTR dynamics with gene expression in the eukaryotic transcriptome. RNA. 2015 Aug;21(8):1502-10. doi: 10.1261/rna.048355.114. Epub 2015 Jun 19. PMID: 26092945; PMCID: PMC4509939.

#This section identifies the input files (.fastq format)
	echo "How do you wish to label your output files (genotype/treatment/etc...)?"
	read genotype
	
	echo "This script assumes you have two replicates of non-paired end illumina sequencing reads in fastq format (not gzipped). Please select the replicate1.fastq file from the list"
		files=$(ls *.{fastq,fq})
		i=1

			for j in $files
				do
					echo "$i.$j"
					file[i]=$j
					i=$(( i + 1 ))
				done
				
			echo "Enter number"
			read input
			echo "You select the file ${file[$input]}"

  	echo "Please select the replicate2.fastq file from the list"
		files=$(ls *.fastq)
		i=1

			for j in $files
				do
					echo "$i.$j"
					file[i]=$j
					i=$(( i + 1 ))
				done
			
			echo "Enter number"
			read input2
			echo "You select the file ${file[$input2]}"	

#This section defines the gene-specific search sequences
	search1=CTCGGTTCATTTTAAATATG
	search2=CATTTTTCTTATTCTAGACT
	search3=TCCTTTTCTTTATAACTTGT
	search4=CATTTAATGAATTGTAATTC
	search5=AGATTAAGAGTATAAGCTTT
	search6=TTCGAATTATTGAAGCTCAA

#This section defines the maximum gap between the 3'end of the search sequence and the beginning of the polyA tail
	gap=10


#This section makes .fasta files from the .fastq files and removes spaces from the record names
	awk 'NR % 4 == 1 || NR % 4 == 2' ${file[$input]} | sed -e 's/@/>/' | sed -e 's/ /_/' > $genotype.R1.fasta
	awk 'NR % 4 == 1 || NR % 4 == 2' ${file[$input2]} | sed -e 's/@/>/' | sed -e 's/ /_/' > $genotype.R2.fasta;
	
#This section makes a table from the fasta files generated in the previous step.
	paste -d "," - - < $genotype.R1.fasta > $genotype.R1.csv
	paste -d "," - - < $genotype.R2.fasta > $genotype.R2.csv
	
#This section performs the search for the pattern based upon the variables input above and outputs a fasta file of hits. Part of the search criteria includes the sequence tags from the polyA test sequencing protocol (GATCGGAA; Harrison PF, Powell DR, Clancy JL, Preiss T, Boag PR, Traven A, Seemann T, Beilharz TH. PAT-seq: a method to study the integration of 3'-UTR dynamics with gene expression in the eukaryotic transcriptome. RNA. 2015 Aug;21(8):1502-10. doi: 10.1261/rna.048355.114. Epub 2015 Jun 19. PMID: 26092945; PMCID: PMC4509939.) If you change the identity of the sequence tag and it's length, you will need to compensate in the 3rd line, replacing "8" with the length of the sequence tag selected.

		ugrep --tag "$search1.{0,$gap}AAAA+GATCGGAA" $genotype.R1.csv > hits0.$genotype.$search1.csv
		ugrep --tag "$search1.{0,$gap}AAAA+GATCGGAA" $genotype.R2.csv >> hits0.$genotype.$search1.csv
		ugrep --tag -Z1 "$search1.{0,$gap}AAAA+GATCGGAA" $genotype.R1.csv > hits1.$genotype.$search1.csv
		ugrep --tag -Z1 "$search1.{0,$gap}AAAA+GATCGGAA" $genotype.R2.csv >> hits1.$genotype.$search1.csv
		ugrep --tag -Z2 "$search1.{0,$gap}AAAA+GATCGGAA" $genotype.R1.csv > hits2.$genotype.$search1.csv
		ugrep --tag -Z2 "$search1.{0,$gap}AAAA+GATCGGAA" $genotype.R2.csv >> hits2.$genotype.$search1.csv
		ugrep --tag -Z3 "$search1.{0,$gap}AAAA+GATCGGAA" $genotype.R1.csv > hits3.$genotype.$search1.csv
		ugrep --tag -Z3 "$search1.{0,$gap}AAAA+GATCGGAA" $genotype.R2.csv >> hits3.$genotype.$search1.csv
	
		ugrep --tag "$search2.{0,$gap}AAAA+GATCGGAA" $genotype.R1.csv > hits0.$genotype.$search2.csv
		ugrep --tag "$search2.{0,$gap}AAAA+GATCGGAA" $genotype.R2.csv >> hits0.$genotype.$search2.csv
		ugrep --tag -Z1 "$search2.{0,$gap}AAAA+GATCGGAA" $genotype.R1.csv > hits1.$genotype.$search2.csv
		ugrep --tag -Z1 "$search2.{0,$gap}AAAA+GATCGGAA" $genotype.R2.csv >> hits1.$genotype.$search2.csv
		ugrep --tag -Z2 "$search2.{0,$gap}AAAA+GATCGGAA" $genotype.R1.csv > hits2.$genotype.$search2.csv
		ugrep --tag -Z2 "$search2.{0,$gap}AAAA+GATCGGAA" $genotype.R2.csv >> hits2.$genotype.$search2.csv
		ugrep --tag -Z3 "$search2.{0,$gap}AAAA+GATCGGAA" $genotype.R1.csv > hits3.$genotype.$search2.csv
		ugrep --tag -Z3 "$search2.{0,$gap}AAAA+GATCGGAA" $genotype.R2.csv >> hits3.$genotype.$search2.csv
		
		ugrep --tag "$search3.{0,$gap}AAAA+GATCGGAA" $genotype.R1.csv > hits0.$genotype.$search3.csv
		ugrep --tag "$search3.{0,$gap}AAAA+GATCGGAA" $genotype.R2.csv >> hits0.$genotype.$search3.csv
		ugrep --tag -Z1 "$search3.{0,$gap}AAAA+GATCGGAA" $genotype.R1.csv > hits1.$genotype.$search3.csv
		ugrep --tag -Z1 "$search3.{0,$gap}AAAA+GATCGGAA" $genotype.R2.csv >> hits1.$genotype.$search3.csv
		ugrep --tag -Z2 "$search3.{0,$gap}AAAA+GATCGGAA" $genotype.R1.csv > hits2.$genotype.$search3.csv
		ugrep --tag -Z2 "$search3.{0,$gap}AAAA+GATCGGAA" $genotype.R2.csv >> hits2.$genotype.$search3.csv
		ugrep --tag -Z3 "$search3.{0,$gap}AAAA+GATCGGAA" $genotype.R1.csv > hits3.$genotype.$search3.csv
		ugrep --tag -Z3 "$search3.{0,$gap}AAAA+GATCGGAA" $genotype.R2.csv >> hits3.$genotype.$search3.csv
		
		ugrep --tag "$search4.{0,$gap}AAAA+GATCGGAA" $genotype.R1.csv > hits0.$genotype.$search4.csv
		ugrep --tag "$search4.{0,$gap}AAAA+GATCGGAA" $genotype.R2.csv >> hits0.$genotype.$search4.csv
		ugrep --tag -Z1 "$search4.{0,$gap}AAAA+GATCGGAA" $genotype.R1.csv > hits1.$genotype.$search4.csv
		ugrep --tag -Z1 "$search4.{0,$gap}AAAA+GATCGGAA" $genotype.R2.csv >> hits1.$genotype.$search4.csv
		ugrep --tag -Z2 "$search4.{0,$gap}AAAA+GATCGGAA" $genotype.R1.csv > hits2.$genotype.$search4.csv
		ugrep --tag -Z2 "$search4.{0,$gap}AAAA+GATCGGAA" $genotype.R2.csv >> hits2.$genotype.$search4.csv
		ugrep --tag -Z3 "$search4.{0,$gap}AAAA+GATCGGAA" $genotype.R1.csv > hits3.$genotype.$search4.csv
		ugrep --tag -Z3 "$search4.{0,$gap}AAAA+GATCGGAA" $genotype.R2.csv >> hits3.$genotype.$search4.csv
		
		ugrep --tag "$search5.{0,$gap}AAAA+GATCGGAA" $genotype.R1.csv > hits0.$genotype.$search5.csv
		ugrep --tag "$search5.{0,$gap}AAAA+GATCGGAA" $genotype.R2.csv >> hits0.$genotype.$search5.csv
		ugrep --tag -Z1 "$search5.{0,$gap}AAAA+GATCGGAA" $genotype.R1.csv > hits1.$genotype.$search5.csv
		ugrep --tag -Z1 "$search5.{0,$gap}AAAA+GATCGGAA" $genotype.R2.csv >> hits1.$genotype.$search5.csv
		ugrep --tag -Z2 "$search5.{0,$gap}AAAA+GATCGGAA" $genotype.R1.csv > hits2.$genotype.$search5.csv
		ugrep --tag -Z2 "$search5.{0,$gap}AAAA+GATCGGAA" $genotype.R2.csv >> hits2.$genotype.$search5.csv
		ugrep --tag -Z3 "$search5.{0,$gap}AAAA+GATCGGAA" $genotype.R1.csv > hits3.$genotype.$search5.csv
		ugrep --tag -Z3 "$search5.{0,$gap}AAAA+GATCGGAA" $genotype.R2.csv >> hits3.$genotype.$search5.csv
		
		ugrep --tag "$search6.{0,$gap}AAAA+GATCGGAA" $genotype.R1.csv > hits0.$genotype.$search6.csv
		ugrep --tag "$search6.{0,$gap}AAAA+GATCGGAA" $genotype.R2.csv >> hits0.$genotype.$search6.csv
		ugrep --tag -Z1 "$search6.{0,$gap}AAAA+GATCGGAA" $genotype.R1.csv > hits1.$genotype.$search6.csv
		ugrep --tag -Z1 "$search6.{0,$gap}AAAA+GATCGGAA" $genotype.R2.csv >> hits1.$genotype.$search6.csv
		ugrep --tag -Z2 "$search6.{0,$gap}AAAA+GATCGGAA" $genotype.R1.csv > hits2.$genotype.$search6.csv
		ugrep --tag -Z2 "$search6.{0,$gap}AAAA+GATCGGAA" $genotype.R2.csv >> hits2.$genotype.$search6.csv
		ugrep --tag -Z3 "$search6.{0,$gap}AAAA+GATCGGAA" $genotype.R1.csv > hits3.$genotype.$search6.csv
		ugrep --tag -Z3 "$search6.{0,$gap}AAAA+GATCGGAA" $genotype.R2.csv >> hits3.$genotype.$search6.csv
	

#This section makes the output tables
		ugrep -Eo "AAAA+GATCGGAA" hits0.$genotype.$search1.csv | awk '{print $1","length($1)-8}' > $genotype.0.output.$search1.csv
		ugrep -Eo -Z1 "AAAA+GATCGGAA" hits1.$genotype.$search1.csv | awk '{print $1","length($1)-8}' > $genotype.1.output.$search1.csv
		ugrep -Eo -Z2 "AAAA+GATCGGAA" hits2.$genotype.$search1.csv | awk '{print $1","length($1)-8}' > $genotype.2.output.$search1.csv
		ugrep -Eo -Z3 "AAAA+GATCGGAA" hits3.$genotype.$search1.csv | awk '{print $1","length($1)-8}' > $genotype.3.output.$search1.csv
		
		ugrep -Eo "AAAA+GATCGGAA" hits0.$genotype.$search2.csv | awk '{print $1","length($1)-8}' > $genotype.0.output.$search2.csv
		ugrep -Eo -Z1 "AAAA+GATCGGAA" hits1.$genotype.$search2.csv | awk '{print $1","length($1)-8}' > $genotype.1.output.$search2.csv
		ugrep -Eo -Z2 "AAAA+GATCGGAA" hits2.$genotype.$search2.csv | awk '{print $1","length($1)-8}' > $genotype.2.output.$search2.csv
		ugrep -Eo -Z3 "AAAA+GATCGGAA" hits3.$genotype.$search2.csv | awk '{print $1","length($1)-8}' > $genotype.3.output.$search2.csv
		
		ugrep -Eo "AAAA+GATCGGAA" hits0.$genotype.$search3.csv | awk '{print $1","length($1)-8}' > $genotype.0.output.$search3.csv
		ugrep -Eo -Z1 "AAAA+GATCGGAA" hits1.$genotype.$search3.csv | awk '{print $1","length($1)-8}' > $genotype.1.output.$search3.csv
		ugrep -Eo -Z2 "AAAA+GATCGGAA" hits2.$genotype.$search3.csv | awk '{print $1","length($1)-8}' > $genotype.2.output.$search3.csv
		ugrep -Eo -Z3 "AAAA+GATCGGAA" hits3.$genotype.$search3.csv | awk '{print $1","length($1)-8}' > $genotype.3.output.$search3.csv
		
		ugrep -Eo "AAAA+GATCGGAA" hits0.$genotype.$search4.csv | awk '{print $1","length($1)-8}' > $genotype.0.output.$search4.csv
		ugrep -Eo -Z1 "AAAA+GATCGGAA" hits1.$genotype.$search4.csv | awk '{print $1","length($1)-8}' > $genotype.1.output.$search4.csv
		ugrep -Eo -Z2 "AAAA+GATCGGAA" hits2.$genotype.$search4.csv | awk '{print $1","length($1)-8}' > $genotype.2.output.$search4.csv
		ugrep -Eo -Z3 "AAAA+GATCGGAA" hits3.$genotype.$search4.csv | awk '{print $1","length($1)-8}' > $genotype.3.output.$search4.csv
		
		ugrep -Eo "AAAA+GATCGGAA" hits0.$genotype.$search5.csv | awk '{print $1","length($1)-8}' > $genotype.0.output.$search5.csv
		ugrep -Eo -Z1 "AAAA+GATCGGAA" hits1.$genotype.$search5.csv | awk '{print $1","length($1)-8}' > $genotype.1.output.$search5.csv
		ugrep -Eo -Z2 "AAAA+GATCGGAA" hits2.$genotype.$search5.csv | awk '{print $1","length($1)-8}' > $genotype.2.output.$search5.csv
		ugrep -Eo -Z3 "AAAA+GATCGGAA" hits3.$genotype.$search5.csv | awk '{print $1","length($1)-8}' > $genotype.3.output.$search5.csv
		
		ugrep -Eo "AAAA+GATCGGAA" hits0.$genotype.$search6.csv | awk '{print $1","length($1)-8}' > $genotype.0.output.$search6.csv
		ugrep -Eo -Z1 "AAAA+GATCGGAA" hits1.$genotype.$search6.csv | awk '{print $1","length($1)-8}' > $genotype.1.output.$search6.csv
		ugrep -Eo -Z2 "AAAA+GATCGGAA" hits2.$genotype.$search6.csv | awk '{print $1","length($1)-8}' > $genotype.2.output.$search6.csv
		ugrep -Eo -Z3 "AAAA+GATCGGAA" hits3.$genotype.$search6.csv | awk '{print $1","length($1)-8}' > $genotype.3.output.$search6.csv

#This section makes files for subsequent alignment and logo production
	cat hits0.$genotype.$search1.csv | tr "," "\n" | sed 's/\___/\n/g' | awk 'NR%2==1{print}' > prealigned0.$genotype.$search1.fa
	cat hits1.$genotype.$search1.csv | tr "," "\n" | sed 's/\___/\n/g' | awk 'NR%2==1{print}' > prealigned1.$genotype.$search1.fa
	cat hits2.$genotype.$search1.csv | tr "," "\n" | sed 's/\___/\n/g' | awk 'NR%2==1{print}' > prealigned2.$genotype.$search1.fa
	cat hits3.$genotype.$search1.csv | tr "," "\n" | sed 's/\___/\n/g' | awk 'NR%2==1{print}' > prealigned3.$genotype.$search1.fa
	
	cat hits0.$genotype.$search2.csv | tr "," "\n" | sed 's/\___/\n/g' | awk 'NR%2==1{print}' > prealigned0.$genotype.$search2.fa
	cat hits1.$genotype.$search2.csv | tr "," "\n" | sed 's/\___/\n/g' | awk 'NR%2==1{print}' > prealigned1.$genotype.$search2.fa
	cat hits2.$genotype.$search2.csv | tr "," "\n" | sed 's/\___/\n/g' | awk 'NR%2==1{print}' > prealigned2.$genotype.$search2.fa
	cat hits3.$genotype.$search2.csv | tr "," "\n" | sed 's/\___/\n/g' | awk 'NR%2==1{print}' > prealigned3.$genotype.$search2.fa
	
	cat hits0.$genotype.$search3.csv | tr "," "\n" | sed 's/\___/\n/g' | awk 'NR%2==1{print}' > prealigned0.$genotype.$search3.fa
	cat hits1.$genotype.$search3.csv | tr "," "\n" | sed 's/\___/\n/g' | awk 'NR%2==1{print}' > prealigned1.$genotype.$search3.fa
	cat hits2.$genotype.$search3.csv | tr "," "\n" | sed 's/\___/\n/g' | awk 'NR%2==1{print}' > prealigned2.$genotype.$search3.fa
	cat hits3.$genotype.$search3.csv | tr "," "\n" | sed 's/\___/\n/g' | awk 'NR%2==1{print}' > prealigned3.$genotype.$search3.fa
	
	cat hits0.$genotype.$search4.csv | tr "," "\n" | sed 's/\___/\n/g' | awk 'NR%2==1{print}' > prealigned0.$genotype.$search4.fa
	cat hits1.$genotype.$search4.csv | tr "," "\n" | sed 's/\___/\n/g' | awk 'NR%2==1{print}' > prealigned1.$genotype.$search4.fa
	cat hits2.$genotype.$search4.csv | tr "," "\n" | sed 's/\___/\n/g' | awk 'NR%2==1{print}' > prealigned2.$genotype.$search4.fa
	cat hits3.$genotype.$search4.csv | tr "," "\n" | sed 's/\___/\n/g' | awk 'NR%2==1{print}' > prealigned3.$genotype.$search4.fa
	
	cat hits0.$genotype.$search5.csv | tr "," "\n" | sed 's/\___/\n/g' | awk 'NR%2==1{print}' > prealigned0.$genotype.$search5.fa
	cat hits1.$genotype.$search5.csv | tr "," "\n" | sed 's/\___/\n/g' | awk 'NR%2==1{print}' > prealigned1.$genotype.$search5.fa
	cat hits2.$genotype.$search5.csv | tr "," "\n" | sed 's/\___/\n/g' | awk 'NR%2==1{print}' > prealigned2.$genotype.$search5.fa
	cat hits3.$genotype.$search5.csv | tr "," "\n" | sed 's/\___/\n/g' | awk 'NR%2==1{print}' > prealigned3.$genotype.$search5.fa
	
	cat hits0.$genotype.$search6.csv | tr "," "\n" | sed 's/\___/\n/g' | awk 'NR%2==1{print}' > prealigned0.$genotype.$search6.fa
	cat hits1.$genotype.$search6.csv | tr "," "\n" | sed 's/\___/\n/g' | awk 'NR%2==1{print}' > prealigned1.$genotype.$search6.fa
	cat hits2.$genotype.$search6.csv | tr "," "\n" | sed 's/\___/\n/g' | awk 'NR%2==1{print}' > prealigned2.$genotype.$search6.fa
	cat hits3.$genotype.$search6.csv | tr "," "\n" | sed 's/\___/\n/g' | awk 'NR%2==1{print}' > prealigned3.$genotype.$search6.fa
			
#This section collects some useful information into a file and gets rid of temporary / unecessary files.

printf '%s\n' "Total Reads (replicate 1): $(grep -c ">" $genotype.R1.fasta)" \
"Total Reads (replicate 2): $(grep -c ">" $genotype.R2.fasta)" \
"------------------------------------" \
"$search1" \
"0 mismatch: $(grep -c ">" hits0.$genotype.$search1.csv)" \
"1 mismatch: $(grep -c ">" hits1.$genotype.$search1.csv)" \
"2 mismatch: $(grep -c ">" hits2.$genotype.$search1.csv)" \
"3 mismatch: $(grep -c ">" hits3.$genotype.$search1.csv)" \
"------------------------------------" \
"$search2" \
"0 mismatch: $(grep -c ">" hits0.$genotype.$search2.csv)" \
"1 mismatch: $(grep -c ">" hits1.$genotype.$search2.csv)" \
"2 mismatch: $(grep -c ">" hits2.$genotype.$search2.csv)" \
"3 mismatch: $(grep -c ">" hits3.$genotype.$search2.csv)" \
"------------------------------------" \
"$search3" \
"0 mismatch: $(grep -c ">" hits0.$genotype.$search3.csv)" \
"1 mismatch: $(grep -c ">" hits1.$genotype.$search3.csv)" \
"2 mismatch: $(grep -c ">" hits2.$genotype.$search3.csv)" \
"3 mismatch: $(grep -c ">" hits3.$genotype.$search3.csv)" \
"------------------------------------" \
"$search4" \
"0 mismatch: $(grep -c ">" hits0.$genotype.$search4.csv)" \
"1 mismatch: $(grep -c ">" hits1.$genotype.$search4.csv)" \
"2 mismatch: $(grep -c ">" hits2.$genotype.$search4.csv)" \
"3 mismatch: $(grep -c ">" hits3.$genotype.$search4.csv)" \
"------------------------------------" \
"$search5" \
"0 mismatch: $(grep -c ">" hits0.$genotype.$search5.csv)" \
"1 mismatch: $(grep -c ">" hits1.$genotype.$search5.csv)" \
"2 mismatch: $(grep -c ">" hits2.$genotype.$search5.csv)" \
"3 mismatch: $(grep -c ">" hits3.$genotype.$search5.csv)" \
"------------------------------------" \
"$search6" \
"0 mismatch: $(grep -c ">" hits0.$genotype.$search6.csv)" \
"1 mismatch: $(grep -c ">" hits1.$genotype.$search6.csv)" \
"2 mismatch: $(grep -c ">" hits2.$genotype.$search6.csv)" \
"3 mismatch: $(grep -c ">" hits3.$genotype.$search6.csv)" \
"------------------------------------" \
"Your genotype input: $genotype" \
"Your gap selection: $gap" > $genotype.stats.txt
	
		rm -f $genotype.R1.csv
		rm -f $genotype.R2.csv
		rm -f $genotype.R1.fasta
		rm -f $genotype.R2.fasta
	
```
